# Rats rely on airflow cues for self-motion perception

**DOI:** 10.1101/2023.12.19.572298

**Authors:** Lior Polat, Tamar Harpaz, Adam Zaidel

## Abstract

Self-motion perception is a vital skill for all species. It is an inherently multisensory process, that combines inertial (body-based) and relative (with respect to the environment) motion cues. While extensively studied in human and non-human primates, there is currently no paradigm to test self-motion perception in rodents using both inertial and relative self-motion cues. We developed a novel rodent motion simulator using two synchronized robotic arms to generate inertial, relative or combined (inertial and relative) cues of self-motion. Eight rats were trained to perform a task of heading-discrimination, similar to the popular primate paradigm. Strikingly, the rats relied heavily on airflow for relative self-motion perception, with little contribution from optic flow (performance in the dark was almost as good). Relative self-motion (airflow) was perceived with greater reliability vs. inertial. Disrupting airflow (using a fan or windshield) damaged relative, but not inertial, self-motion perception. However, whiskers were not needed for this function. Lastly, the rats integrated relative and inertial self-motion cues in a reliability-based (Bayesian-like) manner. These results implicate airflow as a dominant cue for self-motion perception in rats, and provide a new domain to investigate the neural bases of self-motion perception and multisensory processing in awake behaving rodents.

## INTRODUCTION

Air – the medium in which we and all land animals exist – is in constant contact with our bodies. As we move around in the environment, a resulting flow of air (in the opposite direction to our movement) is felt on our skin and hair. Yet, despite this continual stimulus on the largest interface that we have with the environment, little is known about the contribution of airflow cues to our perception of self-motion in space.

‘Anemotaxis’, the ability to use airflow cues in order to navigate in the environment, is known to be used by insects (Kennedy & Marsh, 1974; Baker et al., 1984; Kaissling, 1997; Meats & Hartland, 1999; Sane et al., 2007; Budick et al., 2007; Suver et al., 2019). In drosophila, specialized brain mechanisms integrate airflow cues with visual motion cues to provide a multimodal map of space (Okubo et al., 2020). Also bats detect and use airflow when flying (Sterbing-D’Angelo et al., 2011; Cynthia, 2012). Yet, much less is known about anemotaxis in non-flying animals. It has been observed that marsh rats in the wild primarily move either upwind or downwind (rarely crosswind) presumably to help them navigate (Schooley & Branch, 2005). Also, in a laboratory experiment, rats trained to navigate to a source of airflow demonstrated the ability to detect and respond to airflow cues (Yu et al., 2016). Moreover, Mugnaini et al. (2023) recently showed rats use their supraorbital whiskers to perceive wind resulting from an object’s motion. Thus, airflow may be an important cue for navigation in rodents (and land animals in general) and warrants further investigation.

Information from airflow may be integrated together with other cues of self-motion to improve perception thereof. Multisensory integration for self-motion perception has been extensively studied in humans and monkeys (Fetsch et al., 2009; Butler et al., 2010; Zaidel et al., 2011; Cullen, 2012; Zaidel et al., 2013, 2015, 2017; Y. Zhang et al., 2018; Gu, 2018; Hou et al., 2019) with a primary focus on the contribution of visual and vestibular cues. Despite evidence that airflow facilitates the feeling of vection (the sensation of movement of one’s body in space) even in humans (Seno et al., 2011; Murata et al., 2014; Kurosawa et al., 2017), the question whether, and to what extent, land animals use and integrate airflow information for self-motion perception has been little studied. With a unique apparatus to provide airflow to the forehead in humans, the Bremmer group recently found that these airflow cues are integrated (sub-optimally) with visual cues of self-motion perception (Rosenblum, Grewe, et al., 2022; Rosenblum, Kreß, et al., 2022). Whether more naturalistic airflow (e.g., that could be sensed across the whole body) would be integrated in line with Bayesian predictions of optimality remains an open question.

Cues for self-motion can be divided into two main types: i) “inertial” motion cues, which result from acceleration of one’s body through space. Because inertial motion is sensed predominantly via the vestibular system (vestibular lesion damages this ability; Gu et al., 2007) inertial motion stimuli are often loosely termed “vestibular”. However, inertial motion may also stimulate additional somatosensory senses (e.g., proprioception). Therefore, in this study we use the more general term “inertial” motion. ii) “Relative” motion cues – as one moves in the environment, entities in the environment appear to move (relatively) in the opposite direction. Relative motion cues are thus inherently ambiguous – the same stimulus can arise from motion of the individual in a static environment or from motion of the environment relative to a static individual. The best-known relative motion cue is visual optic flow, which can generate a percept of self-motion (especially effective in primates), e.g., via virtual-reality glasses, even when one is objectively stationary in the world (Brandt et al., 1973; Berthoz et al., 1975; Warren, 2008). Airflow, like optic flow, is a relative motion cue – it could (ambiguously) reflect motion of the air (wind) or result from one’s own self-motion through the environment (or a combination thereof).

Multisensory integration for self-motion perception has only been studied in primates. Both humans and monkeys integrate inertial (vestibular) and relative (visual) motion cues to improve perceptual reliability (Fetsch et al., 2009; Butler et al., 2010; Zaidel et al., 2011; Cullen, 2012; Zaidel et al., 2013, 2015, 2017; Y. Zhang et al., 2018; Gu, 2018; Hou et al., 2019). It is not known whether rodents (or other non-primates) also integrate multiple self-motion cues in a near-optimal (Bayesian) manner. Rodents have lower visual acuity than primates (Prusky et al., 2000) and rely heavily on other cues, e.g., somatosensory (Kleinfeld et al., 2006; Diamond et al., 2008). Therefore, airflow may provide an important relative motion cue for rodents.

Since there is currently no model to study multisensory integration of inertial and relative motion cues in awake behaving rodents (similar to that used in primates) for this study we developed a novel motion simulator for rats. The motion stimuli were controlled by two industrial robotic arms: one robot held and moved a rat platform (upon which the rat was perched) to generate inertial motion cues; while a second robot held and independently moved a large spherical environment surrounding the rat to control relative motion cues. Different combinations of (de)coupled robot motion generated: i) inertial, ii) relative, or iii) combined (inertial and relative) motion cues. We further dissociated which sensory cues the rats relied on for relative motion perception – visual (light vs. dark conditions) or airflow (by disrupting airflow resulting from relative motion).

Strikingly, the rats relied heavily on airflow cues (with little contribution from optic flow) to discriminate relative motion headings. Moreover, these relative (airflow) motion cues were perceived with greater reliability vs. inertial motion cues. Disrupting airflow cues (using a fan or windshield) damaged this ability. Finally, the rats integrated airflow and inertial motion cues largely according to Bayesian predictions.

## RESULTS

### Novel multisensory motion simulator for rodents

To study multisensory perception of self-motion in awake behaving rats, we built a novel rodent motion simulator with independent control over inertial and relative motion stimuli (Fig. 1A; see also Suppl. Fig. 1 for photographs of the setup). This was done by moving (or not moving) the sphere environment which surrounded the rat (via Robot #2) independently from the motion platform upon which the rat was perched (via Robot #1). ‘Inertial’ motion stimuli were generated (in the dark) by moving the rat platform, and synchronously moving the surrounding sphere (coupled with the platform) to prevent any cues of relative motion (Fig. 1A, top schematic). ‘Relative’ motion stimuli were generated by moving only the sphere environment, while the rat platform remained stationary (no inertial motion; Fig. 1A, middle schematic). ‘Combined’ (inertial and relative) motion stimuli were generated by moving only the rat platform without moving the sphere, thereby allowing relative motion cues to be elicited naturally from the environment (Fig. 1A, bottom schematic).

**Figure 1:**
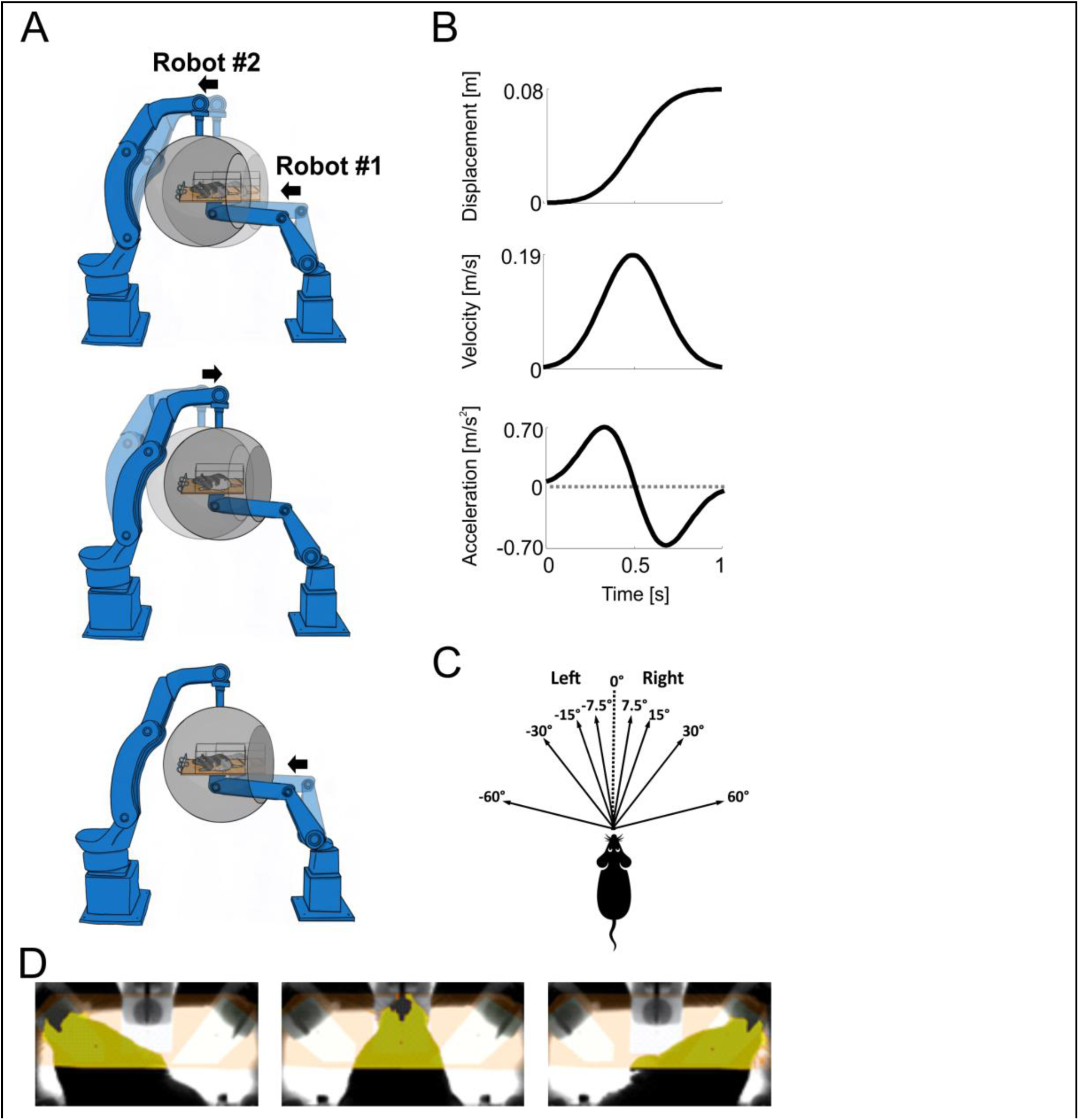
The rat motion system, stimuli and task. (A) Schematics of robot motions (side view) to elicit the self-motion stimuli. Robot #1 supports and moves the rat platform. Robot #2 supports and moves the black sphere (environment). Lighter and darker shades depict the beginning and end positions of the robots, respectively, and arrows adjacent to the robots depict their directions of motion. All three schematics here depict a forward self-motion stimulus (straight-ahead for the rat). Top: inertial-only motion stimulus (relative motion cues are cancelled-out by the synchronous motion of Robot #2 together with Robot #1). Middle: relative-only motion stimulus (only the sphere environment is moved, by Robot #2). Bottom: combined motion stimulus (motion of Robot #1 alone). (B) Dynamic profile of a single self-motion stimulus as a function of time. (C) Schematic of the possible heading directions (viewed from above; 0° marks straight ahead). (D) Infra-red head tracking (from above). Yellow patches show online detection of a rat’s head, drinking from the left, center and right water ports (left, center and right images, respectively). Photographs of the setup are presented in Suppl. Fig. 1.

Eight rats performed a task of heading discrimination. The stimuli comprised linear self-motions in a primarily forward heading direction with deviations (in the horizontal plane) to the right or left of straight-ahead (Fig. 1C). After each self-motion stimulus had ended, the rats were required to report whether their perceived heading was to the right or left of straight-ahead (two-alternative forced choice) by orienting their head to the corresponding side, where they received a liquid reward for correct responses (Fig. 1D). The rats were tested with inertial, relative and combined self-motion cues. To measure cue-integration weights, a heading discrepancy (Δ^+^ = +20° or Δ^−^ = –20°, counterbalanced within each block) was introduced between the inertial and relative heading directions for the combined cues by adjusting the robots’ motions accordingly. The *Baseline* condition thus comprised four stimulus types (interleaved): inertial, relative, and two combined (Δ^+^ and Δ^−^).

### Rats discriminate heading using both inertial and relative self-motion cues

The data for each stimulus type and rat were fit with psychometric curves (cumulative Gaussian; see Methods sub-section **Data acquisition and analysis** for details) that depict the proportion of rightward choices as a function of heading (Fig. 2A). All rats performed better with combined cue stimuli (light and dark green curves) vs. inertial (blue) and relative (red) cues alone. Better performance for the combined cues is seen by two characteristics in their psychometric curves: i) the curves are steeper – meaning, smaller differences in heading could be discriminated (lower perceptual thresholds) and ii) the curve asymptotes are closer to 0 and 1 (for the more eccentric leftward and rightward headings, respectively) – meaning, fewer misses for easy stimuli (lower lapse rates). Both of these factors contribute to better overall performance (compared quantitatively below).

**Figure 2:**
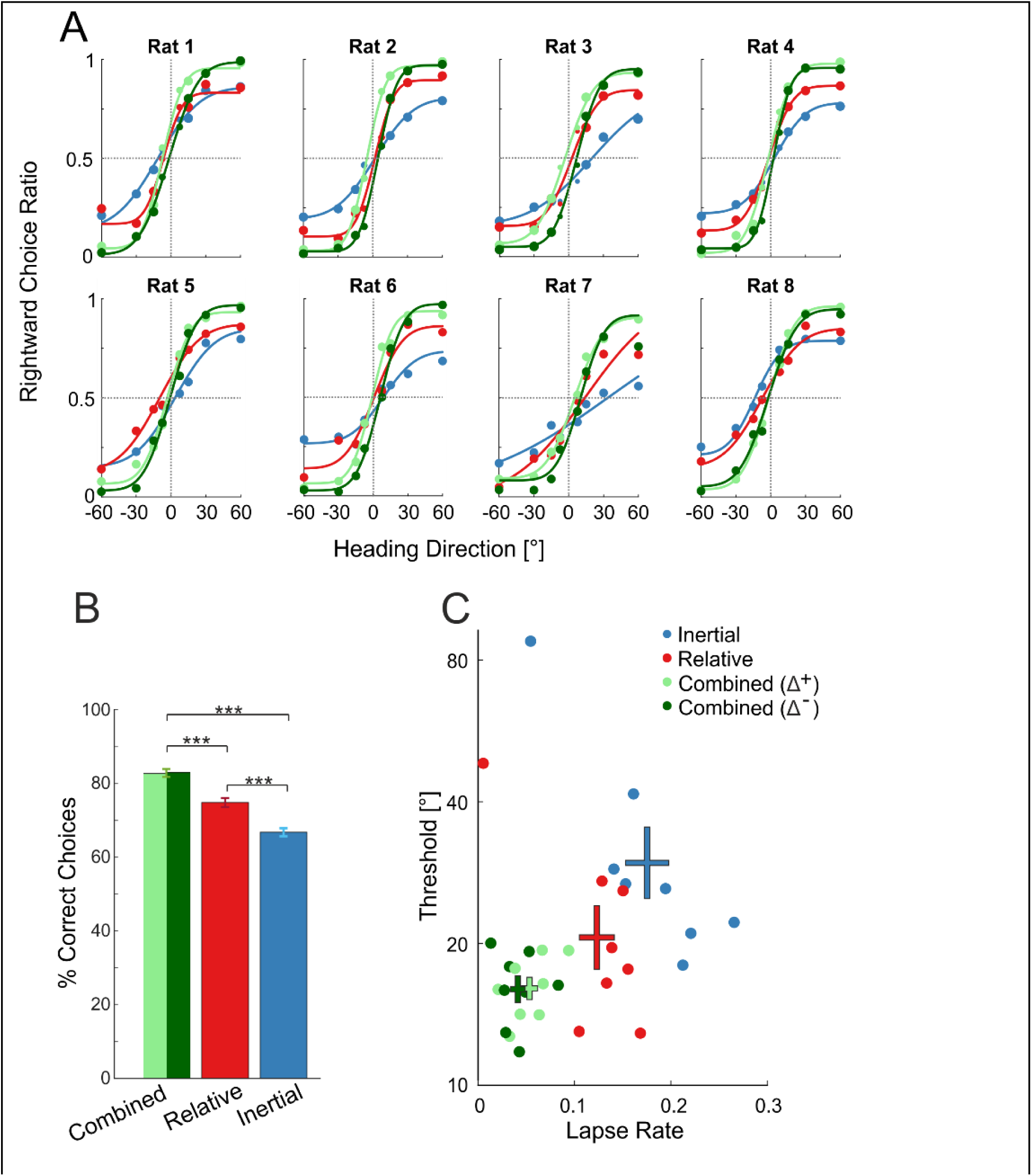
Improved heading discrimination for combined cues. (A) Behavioral responses of eight rats (separate subplots) for: inertial (blue), relative (red) and two combined (light and dark green) stimulus types. The combined stimuli had a systematic offset (Δ) between the inertial and relative motion headings. For Δ^+^ = +20° (light green) relative and inertial motion headings were offset by 10° to the right and left, respectively; and vice versa for Δ^−^ = –20° (dark green). Data points (circles) mark the proportion of rightward choices per heading and stimulus type, and solid curves depict the fitted psychometric functions. Circle sizes reflect the number of trials for each data point. All pseudo-*R²* values were > 0.90. (B) Mean ± SEM percent correct choices per stimulus type. Separate bars (juxtaposed, shades of green) are presented for the two combined stimulus types. The combined cue mean ± SEM was calculated using average values (per rat) from the two combined stimulus types. *** *p* < 0.001. (C) Thresholds and lapse rates from the psychometric fits (per rat and stimulus type). ‘+’ symbols present mean ± SEM values.

First, we compared overall performance between the cues, by looking at the percent correct choices (Fig 2B). Percent correct is a robust (model-free) measure of performance because it does not depend on the psychometric fit. For combined cues, the average percent correct from the two combined stimulus types (Δ^+^ and Δ^−^), per rat, was used for comparison. Percent correct was highest for combined cues (82.8% ± 1.1%; mean ± SEM), followed by relative (74.8% ± 1.2%) and then inertial (66.8% ± 1.1%) cues. These differed significantly across the three cues (*p* = 7.4·10^-9^, *F*_(2,14)_ = 94.6, *η*^2^ = 0.93; repeated measures ANOVA) and each was significantly different from the other two (Holm post-hoc comparisons for repeated measures). Namely, performance was significantly better for combined cues vs. both relative (*p* = 1.5·10^-5^, *t* = 6.87, *Cohen’s d* = 2.43) and inertial (*p* = 4.8·10^-9^, *t* = 13.8, *Cohen’s d* = 4.86), and relative was significantly better vs. inertial (*p* = 1.5·10^-5^, *t* = 6.88, *Cohen’s d* = 2.43).

We further investigated differences in performance by comparing the psychometric fit parameters across cues, specifically: thresholds (quantified by the standard deviation (SD) of the Gaussian fit) and lapse rates (quantified by the psychometric fit asymptotes). We did not compare the point of subjective equality (PSE) because: a) it measures bias, not reliability (Zaidel et al., 2011, 2013, 2021), and b) non-zero values are expected for the combined cue PSEs by design (because of Δ).

A scatter plot of thresholds vs. lapse rates for the different cues, per rat, is presented in Figure 2C. Lower values (for both measures) indicate better performance. Performance on both measures was best for combined cues (green data points lie toward the bottom left), followed by relative (red) and then inertial (blue) cues. Measures from the two combined cue stimulus types (Δ^+^ and Δ^−^) were averaged (per rat) for statistical comparisons across cues. Thresholds differed significantly across combined, relative and inertial cues (mean ± SEM = 15.95° ± 1.05°, 20.69° ± 1.17°, and 29.67° ± 1.19°, respectively; *p* = 0.004, *F*_(2,14)_ = 8.4, *η*^2^ = 0.54, repeated measures ANOVA). Post-hoc comparisons (Holm) indicate that combined cue thresholds were significantly lower vs. inertial cues (*p* = 0.003, *t* = -4.1, *Cohen’s d* = -1.44) and tended to be lower (but not significantly) vs. relative cues (*p* = 0.12, *t* = -1.7, *Cohen’s d* = -0.59). Relative cue thresholds also tended to be lower (but not significantly) vs. inertial (*p* = 0.061, *t* = -2.4, *Cohen’s d* = -0.85).

Lapse rates also differed significantly across combined, relative and inertial cues (mean ± SEM = 5% ± 1%, 12% ± 2%, and 18% ± 2%, respectively; *p* = 3.2·10^-4^, *F*_(2,14)_ = 15.1, *η*^2^ = 0.68, repeated measures ANOVA). Post-hoc comparisons (Holm) found that each cue was significantly different from the other two. Namely, combined cue lapse rates were significantly lower vs. both inertial cues (*p* = 2.5·10^-4^, *t* = -5.5, *Cohen’s d* = -1.93) and relative cues (*p* = 0.012, *t* = -3.2, *Cohen’s d* = -1.15). Also, relative cue lapse rates were significantly lower vs. inertial (*p* = 0.043, *t* = -2.2, *Cohen’s d* = -0.79).

Thus, performance was best for the combined cue, followed by the relative, and then the inertial cues. This is reflected both in thresholds and in lapse rates, leading to a large (compounded) difference in percent correct. Better performance in the combined cue condition presumably reflects the multisensory benefit of cue integration. Before delving deeper into the multisensory effect, we first investigated what sensory information the rats used for perceiving relative self-motion stimuli (which, perhaps surprisingly, had better performance vs. inertial cues). For this we ran a series of control conditions (presented below).

### Sphere removed – relative cue performance dropped to chance level

To verify that performance for relative motion cues depended on actual relative motions of the sphere environment (and not for example, noises from the robots), we dismounted the sphere from Robot #2 and reran the experiment (*No Sphere* condition). In this condition, the robots still performed the identical movements as in the *Baseline* condition. In this state, the rats were exposed to the broader environment of the room (which was of course static). Accordingly, during ‘relative’ stimuli, there was no longer relative motion of the environment in relation to the rat. Rather, Robot #2’s arm just moved overhead (without the sphere), while the environment (room) remained static relative to the rat. Thus, there were no actual relative motion cues, but the robot still made the same motions and noises.

In the *No Sphere* condition, the rats’ performance for ‘relative’ motion cues dropped to near chance level (Fig. 3B, red data points mark the mean ± SEM, across rats, per heading). Percent correct for relative cues in this condition was 49.3% ± 0.9% (mean ± SEM, across rats), which was significantly worse than *Baseline* (*p* = 1.6·10^-5^, Bonferroni corrected; further details in Table 1, row 1) and not significantly above 50% chance level (*p* = 0.77 (uncorrected); further details in Table 1, row 2). This confirms that performance in the *Baseline* condition was based on actual relative motion of the sphere environment, and not on other trivial cues, such as robot noise.

**Figure 3:**
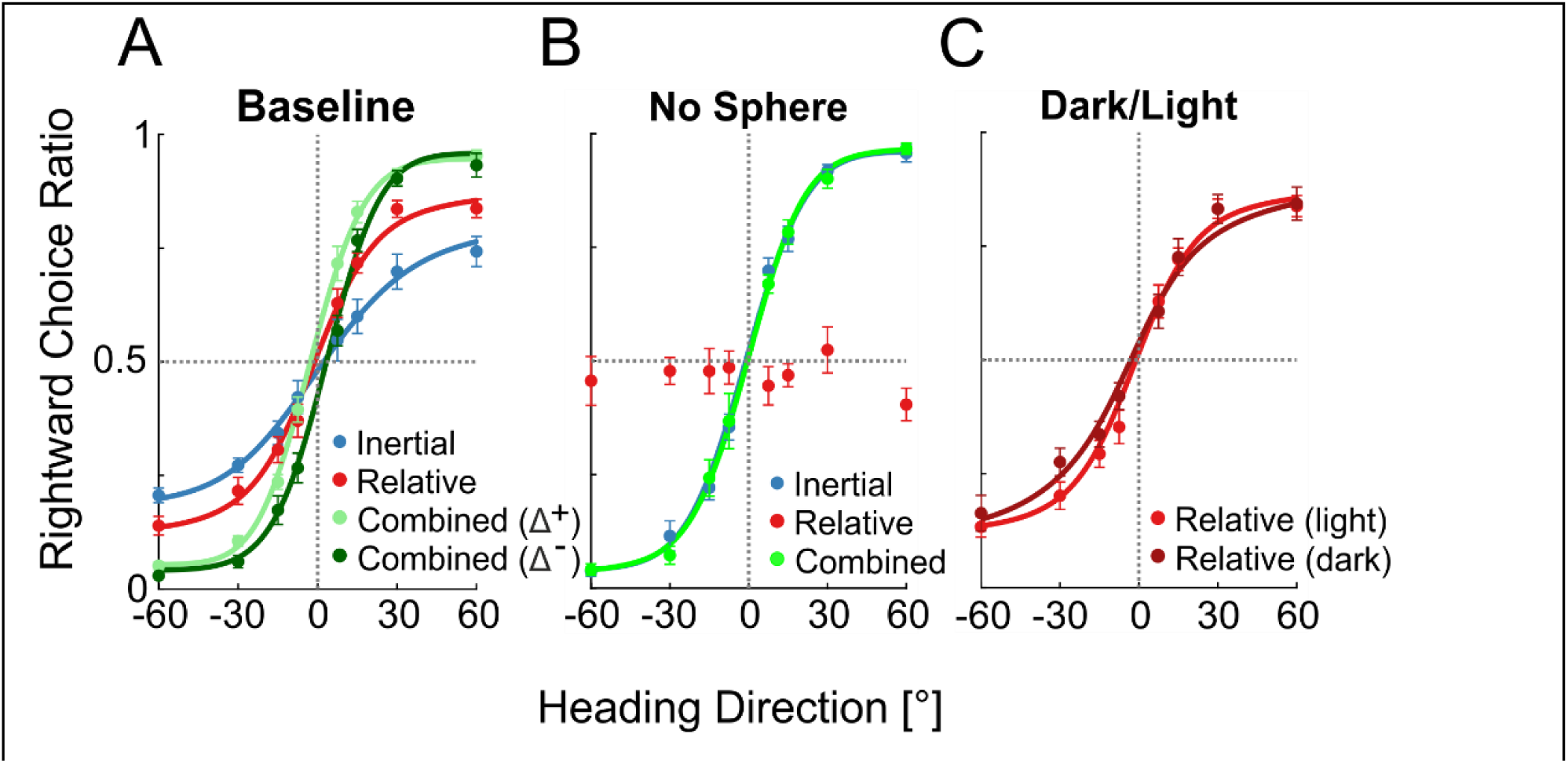
Removing the sphere leads to chance performance for relative motion stimuli. (A) Behavioral performance in the *Baseline* condition (with the sphere mounted on Robot $2, as in Fig. 1 and Suppl. Fig. 1). (B) Behavioral performance in the *No Sphere* condition, with the sphere removed from Robot $2 (but both robots still performing the motions). For the *No Sphere* condition, the combined cue stimulus was run with no cue offset (fluorescent green). (C) Discrimination of relative motion stimuli with LED lights on inside the sphere (‘light’ condition; lighter shade of red in the plot) or off (‘dark’ condition, darker shade of red in the plot). Data points mark the proportion of rightward choices per heading and stimulus (mean ± SEM, across rats) and solid curves depict the fitted psychometric functions, per stimulus. All *pseudo-R^2^* values were > 0.96.

**Table 1:** Statistical comparisons of percent correct for the *No-Sphere* condition

| Row | Stimulus (condition) | Mean ± SEM | vs. | | Comparison Type | $p$ ($p$-Bonf.) | $t$ | Cohen's $d$ | $BF_{10}$ | Conclusion<sup>†</sup> |
| --- | --- | --- | --- | --- | --- | --- | --- | --- | --- | --- |
| 1 | 'Relative' (*No-Sphere*) | 49.3% ± 0.9% | Relative (*Baseline*) | 74.8% ± 1.2% | One-tailed ($H_1$: worse vs. Baseline) | 2.7·10<sup>-6</sup> (1.6·10<sup>-5</sup>) | -16.91 | -6.39 | 5011 | **Worse** |
| 2 | | | Chance | 50% | One-tailed ($H_1$: better vs. chance) | 0.77 (>1) | -0.79 | -0.3 | 0.23 | **Not Better** |
| 3 | 'Inertial' (*No-Sphere*) | 82.7% ± 0.7% | Inertial (*Baseline*) | 66.8% ± 1.1% | Two-tailed | 1.9·10<sup>-5</sup> (9.5·10<sup>-5</sup>) | 12.15 | 4.59 | 1018 | **Better** |
| 4 |  |  | Combined (*No Sphere*) |  | Two-tailed | 0.89 (>1) | 0.15 | 0.06 | 0.35 | Not different (anecdotal evidence) |
| 5 | Combined (*No-Sphere*) | 82.5% ± 1.6% | Combined (*Baseline*) | 82.9% ± 1.1% | Two-tailed | 0.73 (>1) | -0.36 | -0.14 | 0.37 | Not different (anecdotal evidence) |

All comparisons were across 7 rats. Bonferroni corrected $p$ -values are presented in parenthesis (see Methods for further details). <sup>†</sup>Conclusions with substantial evidence ( $BF_{10} > 3$ or $< \frac{1}{3}$ ) are presented in bold, and with anecdotal evidence in regular font. Quotation marks are used for 'relative' and 'inertial' stimulus types in the *No-Sphere* condition because without the sphere they no longer comprise those cues.

This conclusion is also supported by the way that behavior changed in the *No Sphere* condition for the ‘inertial’ stimulus type (in which the robots were coupled and moved synchronously). Here, the ‘inertial’ stimulus type practically became combined stimuli, because the external room now naturally provided relative motion cues (no longer cancelled-out by the sphere). Accordingly, ‘inertial’ performance in the *No Sphere* condition (Fig. 3B, blue, largely obscured by the green curve) was greatly improved vs. *Baseline* (Fig. 3A, blue), showing a significant increase in percent correct (*p* = 9.5·10^-5^, Bonferroni corrected; further details in Table 1, row 3). Moreover, ‘inertial’ performance in this condition was indistinguishable from combined cues (Fig. 3B, overlap of green and blue curves) with no significant difference in percent correct (*p* = 0.89 (uncorrected); further details in Table 1, row 4). The *No Sphere* condition results thus confirm that movement of the sphere environment indeed controls relative self-motion stimuli for the rat.

Moreover, combined cue performance was similar when the relative cues were provided by the external room (*No Sphere* condition) vs. inside the static sphere (*Baseline* condition; *p* = 0.73 (uncorrected); further details in Table 1, row 5). This suggests that performance did not rely on other sphere-specific cues (such as olfaction).

### Minor contribution of optic flow to relative self-motion perception in rats

On trials that contained relative motion stimuli (i.e., relative and combined cue trials) a random combination of LEDs that lined the internal circumference of the sphere were turned on (∼10%, randomly selected on each trial – see Methods sub-section **Experimental conditions** for further details; Suppl. Fig. 1D). Relative motion between the sphere and rat thereby generated optic flow. We initially thought that the rats would rely primarily on optic flow for perceiving relative motion. To test this, we interleaved a fifth stimulus type when running the *Baseline* condition – relative motion in the dark (*Baseline (dark)* condition). In this condition, the LEDs were off (a photometer confirmed that residual light was negligible inside the sphere with LEDs off; see Methods sub-section **Experimental conditions**). We then compared performance for relative cues with LEDs on (light) vs. LEDs off (dark). The rats performed only slightly worse with LEDs off vs. on (dark vs. light red curves in Fig. 3C). Although there was a significant reduction in percent correct (*p* = 0.024, Bonferroni corrected; further details in Table 2A, row 6) the difference was small (72.0% ± 1.8% in dark vs. 74.8% ± 1.2% in light; mean ± SEM). Thus, the benefit of visual cues for perceiving relative stimuli was minor (this was further validated in the other control conditions presented below). This indicates that another (non-visual) sensory modality mostly accounts for perception of relative motion in rats.

**Table 2:** Statistical comparisons of percent correct for control vs. *Baseline* conditions

A) Relative stimulus:
| Row | Condition | Mean ± SEM | $P$<br>($p$-Bonf.) | $t$ | Cohen's $d$ | $BF_{10}$ | Conclusion<sup>†</sup> | vs. |
| --- | --- | --- | --- | --- | --- | --- | --- | --- |
| 1 | *Baseline* | 74.8% ± 1.2% | | | | | | |
| 2 | *Fan* | 59.0% ± 1.2% | $1.8 \cdot 10^{-5}$<br>(0.0001) | -9.21 | -3.26 | 1263 | **Worse** | *Baseline* |
| 3 | *Fan Noise* | 79.1% ± 0.8% | 1.0<br>($>1$) | 5.37 | 2.03 | 0.05 | **Not worse** | |
| 4 | *Windshield* | 61.2% ± 0.8% | $1.5 \cdot 10^{-6}$<br>($9.0 \cdot 10^{-6}$) | -16.61 | -6.28 | 9193 | **Worse** | |
| 5 | *Whiskers Cut* | 75.8% ± 1.0% | 0.73<br>($>1$) | 0.65 | 0.25 | 0.24 | **Not worse** | |
| 6 | *Baseline (dark)* | 72.0% ± 1.8% | 0.004<br>(0.024) | -3.73 | -1.32 | 16.5 | **Worse** | |
| 7 | *Fan (dark)* | 56.6% ± 0.6% | $1.4 \cdot 10^{-4}$<br>(0.0007) | -6.71 | -2.37 | 238 | **Worse** | *Baseline*<br>*(dark)* |
| 8 | *Fan Noise (dark)* | 74.0% ± 0.5% | 0.84<br>($>1$) | 1.08 | 0.41 | 0.20 | **Not worse** | |
| 9 | *Windshield (dark)* | 58.5% ± 0.9% | $2.0 \cdot 10^{-4}$<br>(0.001) | -7.11 | -2.69 | 174 | **Worse** | |
| 10 | *Whiskers Cut (dark)* | 71.9% ± 0.1% | 0.48<br>($>1$) | -0.06 | -0.02 | 0.37 | Not worse<br>(anecdotal evidence) | |

B) Inertial stimulus:
| Row | Condition | Mean ± SEM | $P$<br>($p$-Bonf.) | $t$ | Cohen's $d$ | $BF_{10}$ | Conclusion<sup>†</sup> | vs. |
| --- | --- | --- | --- | --- | --- | --- | --- | --- |
| 1 | *Baseline* | 66.8% ± 1.1% | | | | | | |
| 2 | *Fan* | 64.4% ± 1.3% | 0.13<br>(0.65) | -1.25 | -0.44 | 1.05 | Inconclusive | *Baseline* |
| 3 | *Fan noise* | 67.8% ± 1.6% | 0.69<br>($>1$) | 0.54 | 0.20 | 0.26 | **Not worse** | |
| 4 | *Windshield* | 68.0% ± 1.3% | 0.81<br>($>1$) | 0.95 | 0.36 | 0.21 | **Not worse** | |
| 5 | *Whiskers cut* | 69.6% ± 1.3% | 0.97<br>($>1$) | 2.35 | 0.89 | 0.15 | **Not worse** | |

All comparisons present one-tailed paired $t$ -tests and Bayes Factors ( $H_1$ : control condition worse than the *Baseline* or *Baseline (dark)* condition, see rightmost column) across 7 rats (8 rats for the *Fan* and *Fan (dark)* conditions). Bonferroni corrected $p$ -values are presented in parenthesis (see Methods for further details). <sup>†</sup>Conclusions with substantial evidence ( $BF_{10} > 3$ or $< \frac{1}{3}$ ) are presented in bold, and with inconclusive ( $BF_{10} \sim 1$ ) or anecdotal evidence in regular font.

### Rats rely on airflow cues to perceive relative self-motion

Based on the results above, that the rats perceived relative motion in the dark, but nonetheless required relative motion of the environment (performance dropped to chance level with the sphere removed) we hypothesized that the rats’ perception of relative motion might be mediated by airflow cues. When a closed environment moves (like the sphere here, or a vehicle in motion) the air inside moves with it. Thus, the rats might be sensitive to their relative motion within this medium. To test this, we disrupted airflow cues by placing a fan inside the sphere (*Fan* condition). The fan was fixed above the rat chamber facing forwards, to oppose airflow that would result from relative self-motions forward, and thereby to create turbulence. It was turned on throughout the block. We did not try to manipulate the direction of airflow on a trial-by-trial basis. Rather, we reasoned that if the rats use airflow cues, disruption thereof with a fan would impair that ability, but not impair other motion cues, such as inertial motion or optic flow.

Indeed, heading discrimination of relative motions was severely impaired by the fan (Fig. 4A, top panel), even with optic flow present (LEDs on). Percent correct for relative cues in the *Fan* condition (59.0% ± 1.2%) was significantly worse vs. *Baseline* (*p* = 0.0001, Bonferroni corrected; further details in Table 2A, row 2). By contrast, performance was not significantly worse for inertial cues (Fig. 4A, bottom panel; *p* = 0.13 (uncorrected), further details in Table 2B, row 2). Thus, turbulence selectively damages relative motion cues. Similarly, in the *Fan (dark)* condition (with no optic flow cues, LEDs off; Suppl. Fig. 2A) percent correct for relative cues (56.6% ± 0.6%) was significantly worse vs. *Baseline (dark)* (*p* = 0.0007, Bonferroni corrected; further details in Table 2A, row 7).

**Figure 4:**
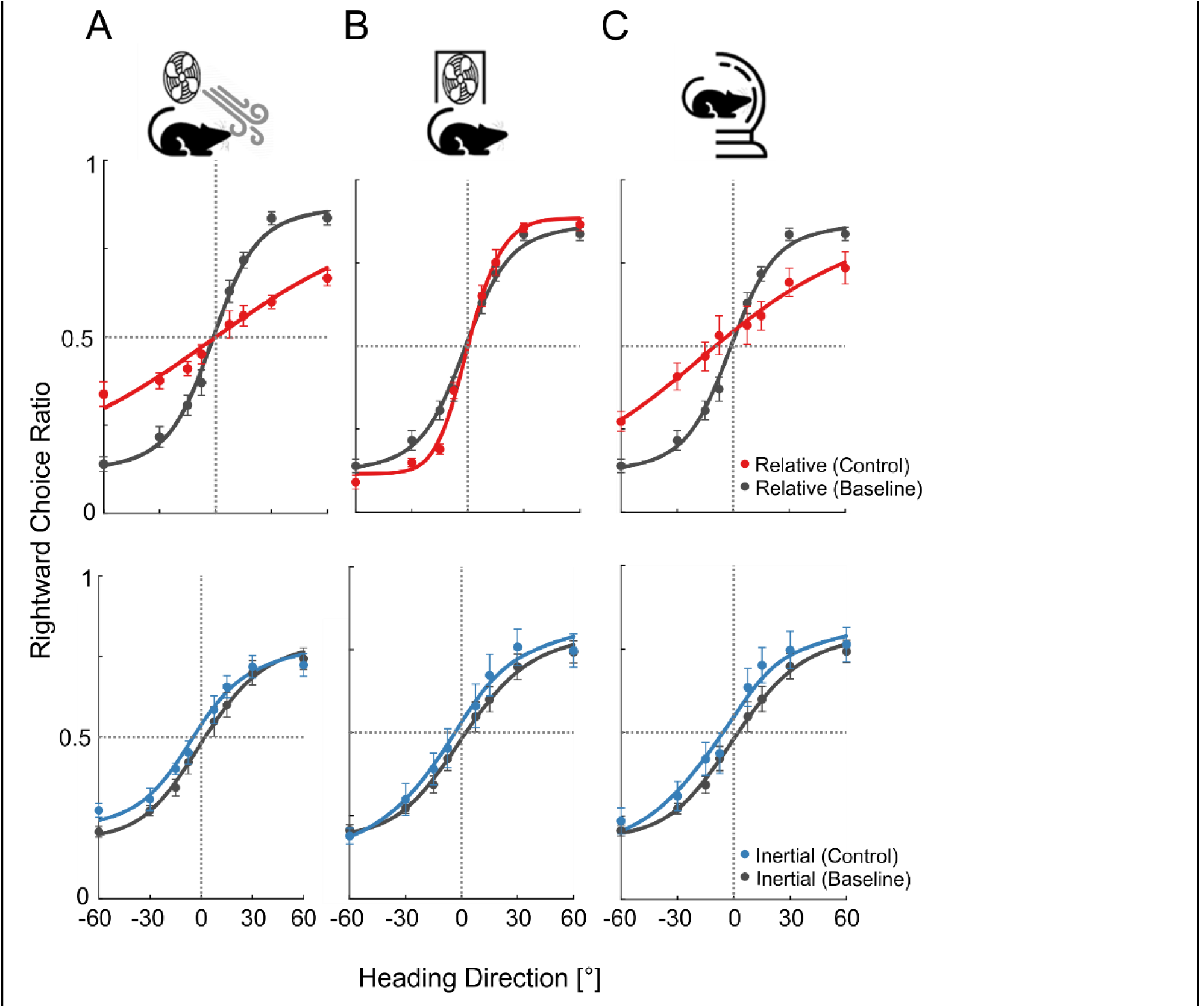
Relative motion discrimination relies on airflow cues. Behavioral performance in three control conditions (columns): (A) Fan: to disrupt airflow cues. (B) *Fan Noise*: the fan was covered to no longer disrupt airflow cues, but still generate fan noise. (C) *Windshield*: to disrupt airflow cues. Psychometric plots for relative and inertial motion stimuli are presented in the top and bottom row, respectively. These are presented in color for the control conditions (red and blue for relative and inertial motion cues, respectively) vs. the Baseline condition (in gray). Data points mark the proportion of rightward choices per heading and stimulus (mean ± SEM, across rats) and solid curves depict the fitted psychometric functions. All pseudo-*R^2^* values were > 0.91. The schematic icons (top) were designed using images from flaticon.com. See also Suppl. Figs. 2 and 3.

It is unlikely that performance was impaired by noise from the fan, because: i) only relative (not inertial) motion perception was worse in the *Fan* condition, and ii) we already validated above (*No Sphere* condition) that the rats do not solve the task by auditory cues from the robots. Nonetheless, to further verify this, we ran the experiment again with the fan, this time covering it with a custom-fit Perspex box to block the air, but maintain fan noise (*Fan Noise* condition; Fig. 4B). No deterioration in percent correct was observed (vs. *Baseline*) for relative cues (*p* = 1.0 (uncorrected); further details in Table 2A, row 3), or inertial cues (*p* = 0.69 (uncorrected); further details in Table 2B, row 3). Similarly, no deterioration was observed for relative cues in the *Fan Noise (dark)* vs. *Baseline (dark)* conditions (*p* = 0.84 (uncorrected); further details in Table 2A, row 8). Thus, air turbulence from the fan (and not fan noise) impaired performance, specifically for relative motion stimuli.

To further validate that rats use airflow for relative self-motion perception, we used a different (passive) way to impair airflow, without an active device. We fixed a windshield made from transparent Perspex to the rat platform, in front of the rats just beyond the reward spouts, to (at least partially) block airflow (*Windshield* condition, Fig. 4C). In this condition, percent correct for relative cues (61.2% ± 0.8%) was significantly worse vs. *Baseline* (*p* = 9.0·10^-6^, Bonferroni corrected; further details in Table 2A, row 4). By contrast, no deterioration was seen for inertial cues (*p* = 0.81 (uncorrected); further details in Table 2B, row 4). Similarly, with LEDs off, in the *Windshield (dark)* condition (Suppl. Fig. 2C) percent correct for relative cues (58.5% ± 0.9%) was significantly worse vs. *Baseline (dark)* (*p* = 0.001, Bonferroni corrected; further details in Table 2A, row 9). Together, these results indicate that the rats used airflow cues for perception of relative self-motion. We next asked whether this was mediated by their whiskers.

### Airflow perception is not mediated by the whiskers

To investigate whether the rats used their whiskers to discriminate airflow cues, we tested performance after trimming all their whiskers (see Methods sub-section **Experimental conditions** for further details). This did not detract from task performance (Suppl. Fig. 3). Performance was not worse in the *Whiskers Cut* condition vs. *Baseline*, neither for relative (*p* = 0.73 (uncorrected); further details in Table 2A, row 5), nor inertial (*p* = 0.97 (uncorrected); further details in Table 2B, row 5) cues. This indicates that the whisker system is not required for discriminating airflow of relative self-motion.

### Summary of performance across conditions

Figure 5 presents a summary plot of performance (percent correct) across conditions. In conditions that did not perturb airflow, performance was consistently better for relative vs. inertial cues (*Baseline*, *Fan Noise*, *Whiskers Cut*; bars with solid texture fill). In conditions that damaged airflow, performance was consistency decreased for relative cues, and became worse vs. inertial cues (*Fan*, *Windshield*, *No Sphere*; bars with diagonal line texture fill). In the *No Sphere* condition, as described above, the ‘relative’ stimulus had no airflow cues while the ‘inertial’ stimulus essentially became combined cues – hence, the extreme decrease and increase in ‘relative’ and ‘inertial’ performance, respectively.

Performance for relative cue stimuli was significantly (and consistently) better with LEDs on (light red bars) vs. LEDs off (dark red bars; *p* = 0.002, *F*_(1,6)_ = 25.14, *η*^2^ = 0.04; repeated measures ANOVA). This indicates a small, but consistent, contribution of optic flow to relative motion perception in rats (secondary to airflow).

### Rats integrate relative and inertial motion cues in a near-optimal Bayesian manner

Lastly, we investigated the multisensory benefit of integrating relative and inertial cues of self-motion in rats. For this, we took advantage of the fact that the different control conditions manipulated relative motion cue reliability, and thereby provided a range of predicted combined (inertial-relative) cue performance for the different conditions. We analyzed this in the Bayesian framework, with quantitative predictions for: i) combined cue thresholds, and ii) cue integration weights (see Methods sub-section **Data acquisition and analysis** for further details).

Figure 6A presents the behavioral thresholds for each cue (inertial, relative and combined) across the different conditions. Inertial thresholds were relatively constant across conditions (blue line; *p* = 0.82, *F*_(4,24)_ = 0.38, *η*^2^ = 0.06; repeated measures ANOVA). By contrast, relative cue performance differed significantly across conditions (red line; *p* = 2.7·10^-11^, *F*_(4,24)_ =50.0, *η*^2^ = 0.89). Specifically, thresholds were significantly higher than *Baseline* in the *Fan* and *Windshield* conditions (*p* = 0.005 and *p* = 0.01, respectively, Bonferroni corrected; further details in Table 3A, rows 2 and 4, respectively). However, not in the *Fan Noise* and *Whiskers Cut* conditions (*p* = 0.99 and *p* = 0.83, respectively (uncorrected); further details in Table 3A, rows 3 and 5, respectively).

**Table 3:**
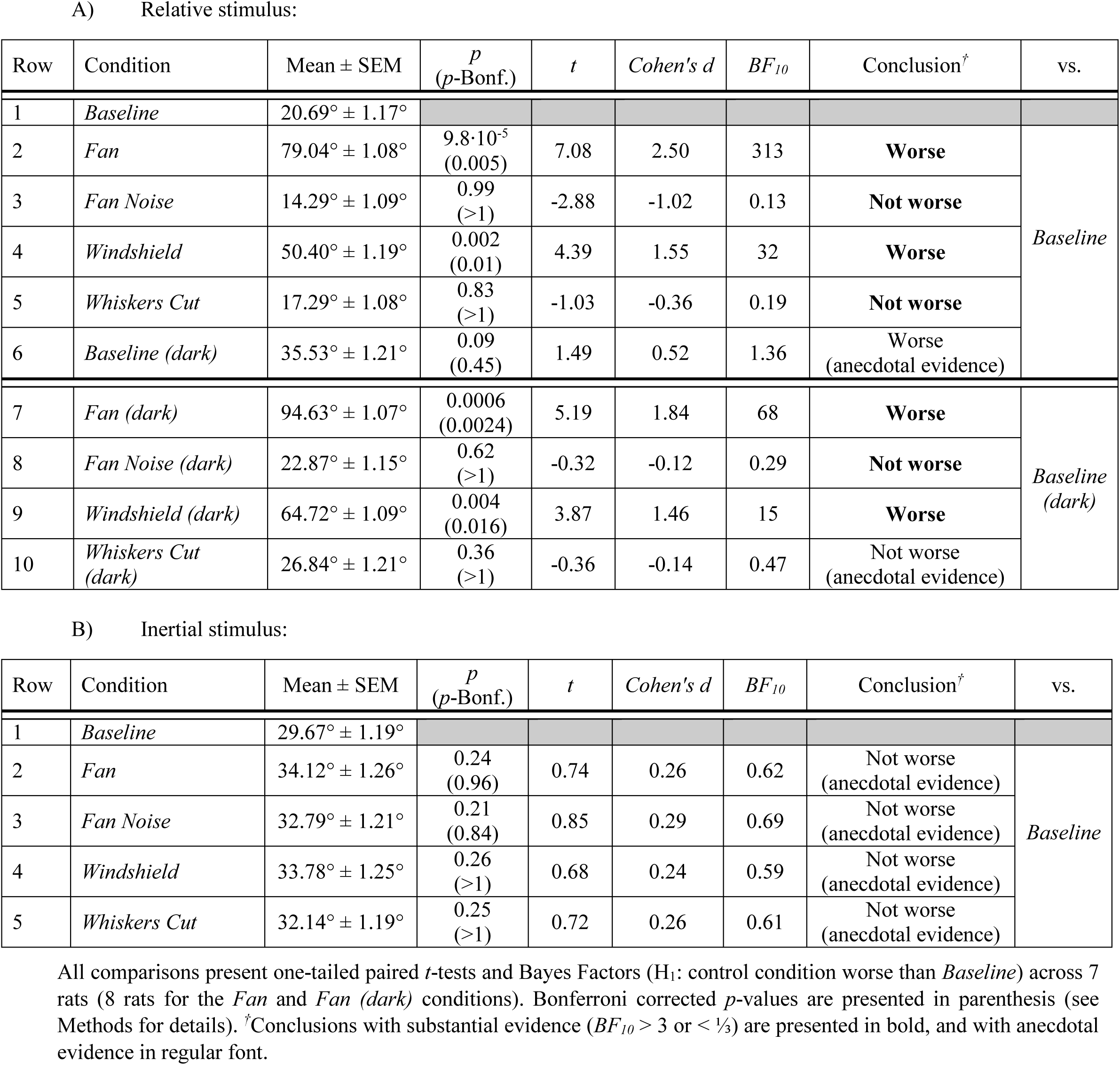
Statistical comparisons of thresholds for control vs. *Baseline* conditions

Combined cue thresholds largely followed Bayesian expectations: when relative thresholds were a lot better (lower) than inertial thresholds (the first three conditions plotted from the left in Fig 6A), combined cue thresholds were slightly better than (or similar to) the relative thresholds. When relative thresholds became larger (worse) than inertial thresholds (two conditions on the right), the combined cue thresholds increased slightly, but remained better than the (now more reliable) inertial cue. To further demonstrate that combined cue thresholds largely followed Bayesian predictions, we plotted the observed vs. Bayesian-predicted combined cue thresholds, per condition (Fig. 6B). The two conditions with increased relative cue thresholds (*Windshield* and *Fan*) also had larger predicted and observed combined cue thresholds (upward and downward facing triangles, respectively). Similarly, the predicted and observed relative cue weights were lower in these two conditions (Fig. 6C). Lastly, when pooling across rats and conditions, the observed and Bayesian predicted values were highly correlated – both for combined cue thresholds (*r*(35) = 0.79, *p* = 1.19·10^-9^, BF_10_ = 1.06·10^7^) and integration weights (*r*(35) = 0.78, *p* = 2.69·10^-9^, BF_10_ =4.95·10^7^, Pearson’s correlations). Thus, the rats reweighted inertial and relative self-motion cues for integration, according to cue reliability.

**Figure 5:**
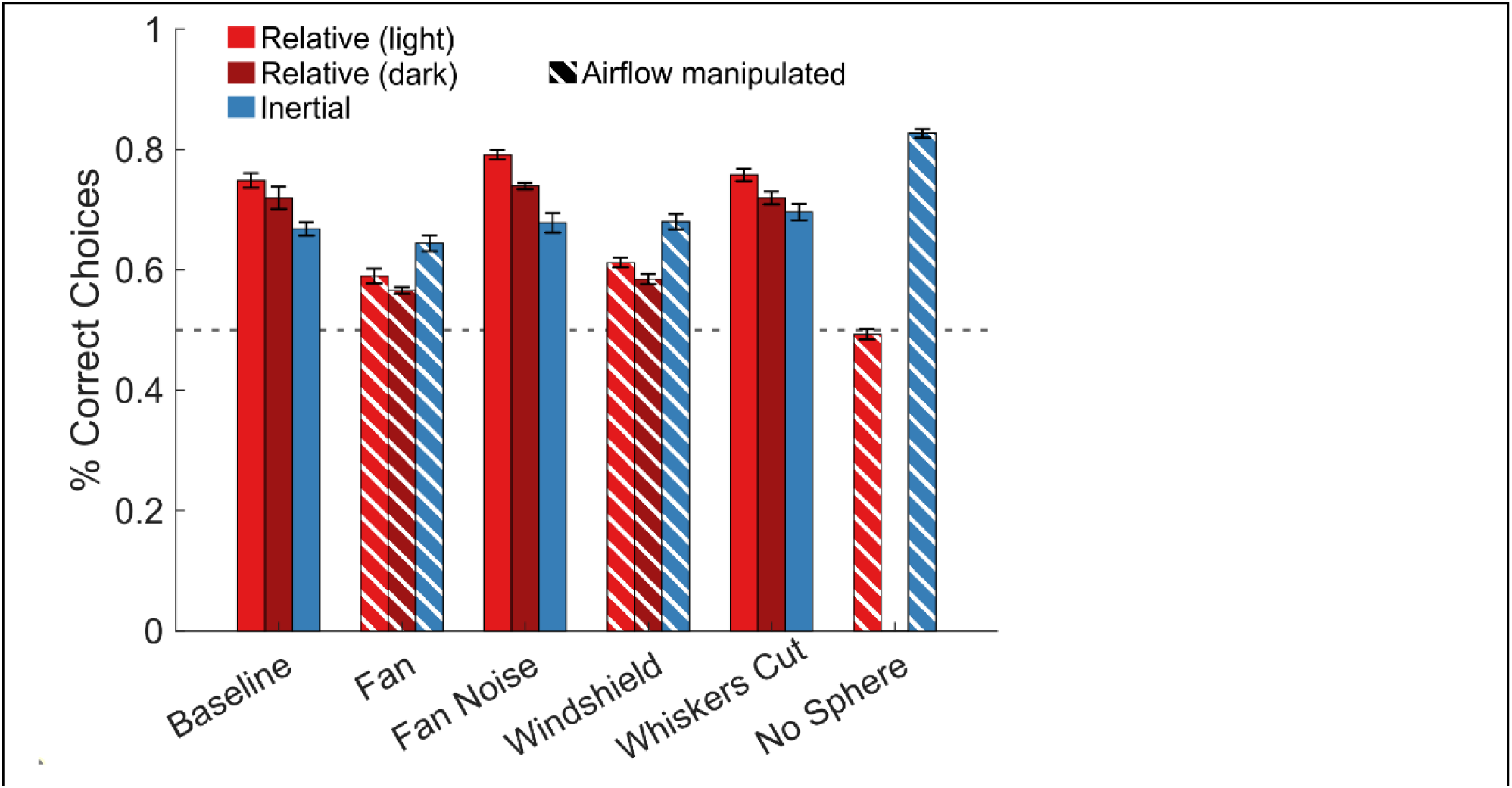
Summary of performance across conditions. Mean ± SEM percent correct choices across rats for relative cues (light and dark shades of red, for LEDs on and off, respectively) and inertial cues (blue). Conditions in which airflow was manipulated are marked by diagonal line texture fill. In the *No Sphere* condition there was no manipulation of LEDs on/off (the larger environment of the room was exposed). The horizontal dashed line marks chance level (0.5).

**Figure 6:**
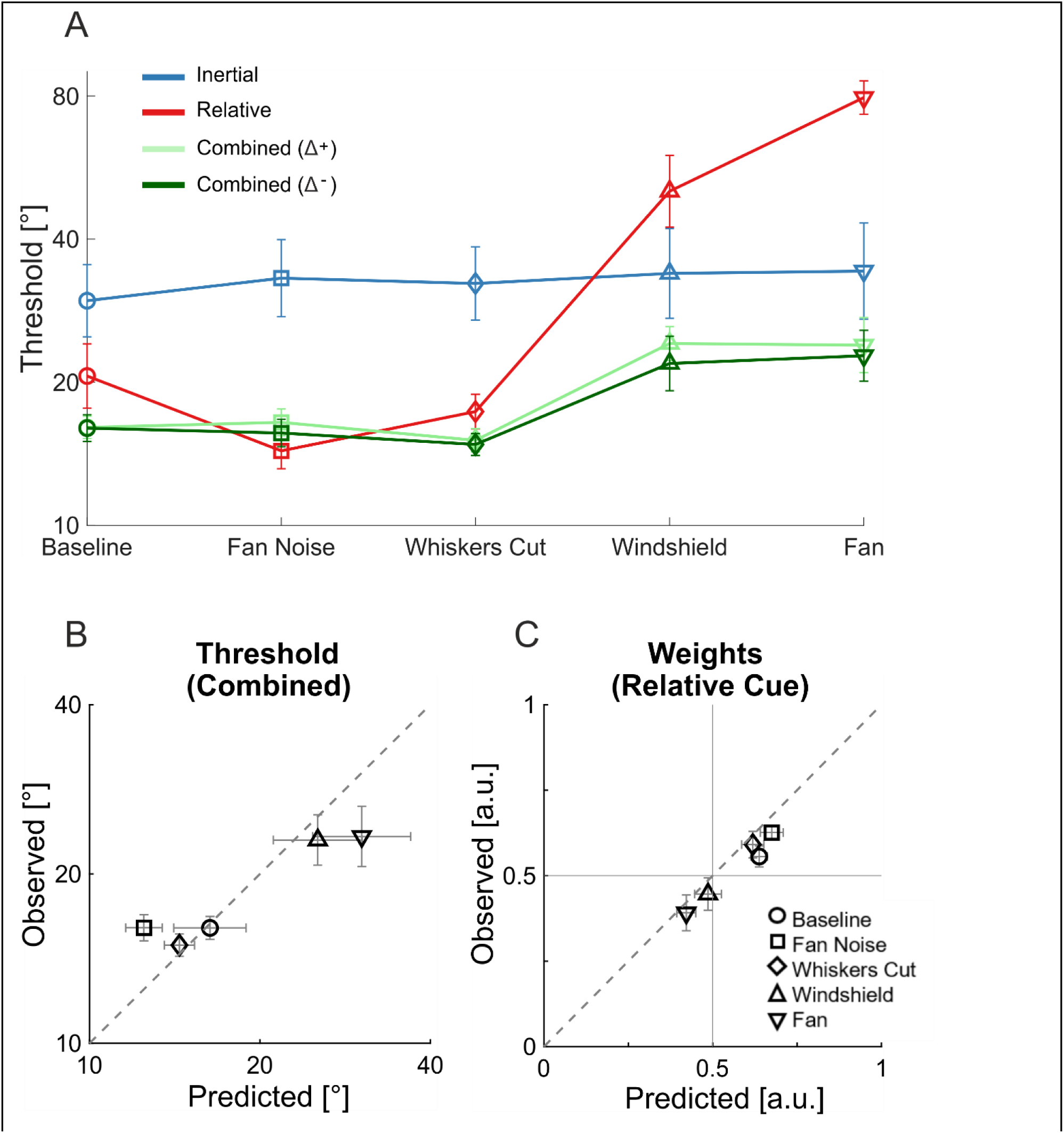
Rats flexibly weight cues according to reliability. (A) Perceptual thresholds (geometric mean ± SEM, across rats) for the different stimulus cues and control conditions. (B) Observed vs. Bayesian-predicted combined cue thresholds (geometric mean ± SEM, across rats) per condition. (C) Observed vs. Bayesian-predicted weights for the relative motion cue (mean ± SEM, across rats) per condition. The horizontal and vertical gray lines in C depict equal cue weighting (0.5). The diagonal dashed lines in B and C mark the line of equality (y = x).

## DISCUSSION

In this study we tested multisensory self-motion perception in awake behaving rats. For this purpose, we developed a novel rodent motion simulator that can separately control inertial and relative motion cues (we believe that it is the first of its kind). We found that rats rely heavily on airflow cues for relative self-motion perception, and that relative (airflow) self-motion cues were perceived with greater reliability vs. inertial self-motion cues. Moreover, rats integrate relative and inertial self-motion cues in a reliability-based (Bayesian like) manner.

### Airflow – a dominant cue for self-motion perception in rats

Initially, we predicted that perception of relative self-motion would be mediated primarily by vision (optic flow). A main, unexpected, finding in this study was that it was rather mediated primarily by airflow cues. We confirmed this through a series of control conditions. Firstly, even in the absence of optic flow, perception of relative self-motion was still highly reliable (and more reliable than inertial self-motion cues). Although optic flow, when present, did improve perceptual reliability of relative self-motion (presumably through integration with airflow cues) its contribution was small. We then removed the sphere environment, but still had the robots perform the same movements. In this state, behavioral performance dropped to chance level for ‘relative’ stimuli. Thus, the rats were not solving the task using some other trivial cue (such as robot noise). Furthermore, disturbing airflow actively (using a fan) or passively (using a windshield) reduced performance, while noise from the fan (without disturbing airflow) did not affect performance. These results confirm that airflow was the dominant cue for self-motion perception in rats.

The finding that rats rely heavily on airflow for relative self-motion perception is in line with recent studies that found that rats spontaneously turn towards airflow stimuli (Mugnaini et al., 2023) and can use airflow cues for navigation (Schooley & Branch, 2005; Yu et al., 2016). Moreover, we found here that airflow is a dominant cue for self-motion perception (more than vision and inertial cues). Together, these results suggest that anemotaxis may be an important (and perhaps underappreciated) function for rodent navigation (Schooley & Branch, 2005).

### Whiskers aren’t used for self-motion perception

It has been previously suggested that whiskers can aid wind sensing in rats. Yu et al. (2016) found decreased localization of airflow in rats after trimming their whiskers. However, they also found that rats without whiskers still performed the airflow localization task well above chance levels, indicating that wind sensing is not entirely whisker based. Mugnaini et al. (2023) specified a particular set of whiskers that have biomechanics which are more sensitive to airflow (the long supra-orbital whiskers, followed by the α, β, and A1 whiskers). They also found that after trimming these whiskers rats turned less toward an airflow stimulus compared to rats who had other (wind-insensitive) whiskers trimmed. Thus, both of these studies describe a reduction in behavioral orienting to wind after whisker trimming. However, in both cases the rats still oriented toward airflow, indicating that whiskers are not the only modality used for airflow sensing. Skin and fur covers a large surface area. Thus it is possible that integration of the many and broad tactile signals felt across the skin (by movement of the fur) can be used for airflow sensing. This is difficult to test (e.g. via inactivation) because of its broad bodily distribution.

In our study, the rats performed the task unhindered, even after whisker trimming. This firstly confirms that airflow perception does not rely entirely on sensing via the whiskers (in line with the two previous studies). Regarding why we found no reduction in performance (unlike the two previous studies), this probably stems from stimulus and task differences. Mugnaini et al. (2023) tested spontaneous turning towards wind stimuli generated by flapping a hand or piece of cardboard. This supposedly reflects a moving object (not self-motion). Yu et al. (2016) trained rats to localize a source of airflow (generated by a fan) and to run toward a hole in front of the fan. Thus, in both cases the rats were required to detect an external source of wind. By contrast, the stimuli in our experiment reflected self-motion only (generated either by moving the rat in the medium or moving the entire medium around the rat). It is possible that whiskers are involved in detecting wind from external objects, but not in self-motion perception. We speculate that self-motion might create a more coherent pattern of airflow across the skin and fur compared to motion of an external object in the environment, which can trigger different patterns of airflow, depending on its source. Accordingly, whiskers might be used for object sensation, but less relevant for self-motion perception.

### Multisensory integration of self-motion cues in rats

Self-motion perception is a vital skill across species – all animals need to know where they are in space in order to behave (the vestibular system is one of the oldest sensory systems phylogenetically (Romer & Parsons, 1978)). It is also an inherently multisensory function. While multisensory integration for self-motion perception has been well-studied in human and non-human primates (Fetsch et al., 2009; Butler et al., 2010; Zaidel et al., 2011; Cullen, 2012; Zaidel et al., 2013, 2015; Y. Zhang et al., 2018; Gu, 2018; Hou et al., 2019; Zaidel et al., 2017), to date, there has been no model to study this in awake behaving rodents.

This novel paradigm of multisensory self-motion perception for rodents can offer many complementary advantages to primates, such as rodent-specific technologies, better affordability and increased sample size. Moreover, it enables cross species comparison to investigate which aspects are preserved across species, and which are species specific (Carandini & Churchland, 2013; Hanks & Summerfield, 2017). Our findings here suggest that airflow is the primary cue for relative self-motion perception in rats, unlike primates (who rely heavily on optic flow). However, we also found similarities – namely, the different cues of self-motion were integrated in a reliability-based manner, similar to primates. This is also in line with other studies that found robust integration of multisensory cues in rats, e.g., audio-visual (Raposo et al., 2012; Sheppard et al., 2013) and visuo-tactile (Nikbakht et al., 2018). Thus, rats offer an excellent model to further investigate the neural bases of self-motion perception and multisensory processing.

### The neural bases

Vestibular inputs project to widespread multisensory cortical areas in both primates (Lopez et al., 2012) and rodents (Rancz et al., 2015). Thus, self-motion perception is a natural model system to study multisensory integration. The neural bases of multisensory integration for self-motion perception has been well studied in monkeys, although there are still many open questions. But, to date, this has not been likewise studied, in awake behaving rodents. In primates, multisensory signals of self-motion are processed in diverse cortical areas. We outline some of these below, and hypothesize based on this (and what is known in rodents) what we might expect to find in rats, in future neuronal studies.

The monkey dorsal medial superior temporal (MSTd) and ventral intraparietal (VIP) areas both have large populations of neurons that respond to visual and vestibular self-motion stimuli (Bremmer et al., 1997, 2002; Chen et al., 2011a; Colby et al., 1993; Duffy, 1998; Gu et al., 2006, 2007; Maciokas & Britten, 2010; Page & Duffy, 2003; Takahashi et al., 2007; T. Zhang et al., 2004; T. Zhang & Britten, 2010). However, MSTd (part of the extra-striate visual cortex) contains more visual signals, while VIP has stronger choice-driven signals (Chen et al., 2013; Zaidel et al., 2017). Closer to the vestibular periphery, parieto-insular vestibular cortex (PIVC) receives stronger and earlier vestibular signals (Chen et al., 2010, 2011a). The visual posterior sylvian area (VPS, strongly interconnected with, and lying posterior to PIVC) also exhibits visual and vestibular self-motion signals (Chen et al., 2011b; Frank et al., 2016). Frontally, the frontal eye fields (FEF, related to oculomotor function) contain strong visual-vestibular self-motion signals (Fukushima et al., 2006; Gu et al., 2016). Thus, there are many multisensory cortical areas, widely distributed across the brain, that encode multisensory self-motion signals. The specific functions of each are not fully understood.

Our findings that airflow is an important cue for self-motion perception in the rat suggest that future work should study cortical areas that can integrate across broad cutaneous fields, e.g., the secondary somatosensory cortex (SSC), which has been previously implicated in multisensory integration (Menzel & Barth, 2005). Also, in accordance with monkey PIVC, vestibular dominant multisensory signals might be found in areas proximal to the rat insular cortex (Rancz et al., 2015). In terms of higher-level decisions, the rat posterior parietal cortex (PPC) is a multimodal association area involved in spatial orientation (McNaughton et al., 1994; Torrealba & Valdés, 2008; Wilber, Clark, Forster, et al., 2014; Wilber et al., 2017), and perceptual decision making (Licata et al., 2017). Thus, multisensory self-motion signals as well as strong decision-related signals may be found in PPC, similar to monkey VIP.

The retrosplenial cortex (RSC) in rats is located medially alongside, and highly interconnected with the PPC (Wilber, Clark, Demecha, et al., 2014). It too is believed to have a key role in spatial orientation and navigation (it has “head direction” cells), as well as higher-level cognitive function (Vann et al., 2009). Both RSC and PPC are also highly (and reciprocally) interconnected with the frontal orienting fields (FOF; known also by other names including M2, Fr2) (Corwin & Reep, 1998; Uylings et al., 2003; Yamawaki et al., 2016), which is believed to be homologous to the primate FEF (Leonard, 1969). Intriguingly, all three areas (PPC, RSC and FOF) have strong vestibular responses (Rancz et al., 2015), receive extensive sensory input (Mohan et al., 2019; Vann et al., 2009; Wilber, Clark, Demecha, et al., 2014) and are involved in orienting movements (Kopec et al., 2015; Mimica et al., 2018) and perceptual decision making (Barthas & Kwan, 2017; Brunton et al., 2013; Licata et al., 2017; Nelson et al., 2014). Therefore, they may support multisensory perception of self-motion. Thus, investigation into the neural basis of self-motion perception in rats (unchartered territory) offers high potential for understanding cross-species fundamentals of multisensory brain function.

### Limitations

We describe here several limitations of our study. Although our control conditions provide substantial evidence that relative self-motion was indeed mediated by airflow, we do not have any biological proof of this. Whisker trimming did not damage performance, and thus, we speculate that this function could be mediated by sensation on the fur and body. However, we did not test this directly. This is difficult to test due to the broad distribution of skin, however, future studies should aim to test this directly. It is possible that neuronal recordings can shed light on this question, if e.g., there is overlap between neuronal responses to somatosensory stimuli on the skin and self-motion stimuli. If so, inactivation of these regions might affect perception of relative self-motion cues.

Regarding our finding that whiskers are not involved in this function – the control conditions were run towards the end of behavioral testing, whereas the *Baseline* condition was run at the beginning. It is therefore possible that performance improved over time, such that the control conditions had a behavioral advantage. While this strengthens findings of impaired function in the control conditions that manipulated airflow (*Fan*, *Windshield*) it could mask deterioration from cutting the whiskers if that deterioration was small. Hence, it is possible that trimming the whiskers did have some (undetected) influence. Nonetheless, our results show that whisker trimming did not have a major effect.

### Future directions

The extent to which airflow can influence perception of self-motion in other species, including humans, has rarely been explored. There is evidence that airflow facilitates the feeling of vection in humans (Seno et al., 2011; Murata et al., 2014; Kurosawa et al., 2017), and Rosenblum et al. (2022) recently showed that airflow toward the forehead in humans influences visual perception of self-motion. Thus airflow does seem to affect self-motion perception in humans. Further investigation, with airflow that can be felt across broader regions of the body, together with vestibular cues, is warranted.

Multisensory processing is a broad and rich field of investigation, which comprises many functions besides multisensory integration. These include multisensory recalibration (Zaidel et al., 2011, 2013, 2021; Zeng et al., 2022; Shalom-Sperber et al., 2022), causal inference (Kording et al., 2007; Acerbi et al., 2018; Dokka et al., 2019) and even perception of one’s self (Zaidel & Salomon, 2023). Moreover, self-motion perception, and other multisensory functions, are often altered in brain disorders such as Parkinson’s disease (Halperin et al., 2020; Yakubovich et al., 2020) and autism (Zaidel et al., 2015; Feigin et al., 2021; Noel et al., 2022). Therefore, this rodent paradigm of self-motion perception provides a basis to further investigate a host of multisensory questions, both behaviorally and neuronally, in health and disease.

## METHODS

### Animal subjects

All experiment procedures were approved by the institutional animal ethics committee at Bar-Ilan University (No. 60-08-2018). Eight male Long-Evans rats were used in this study. For task training and testing, a water restriction protocol was used, by which the rats were rewarded in the experiment for completing trials, and for making correct choices (details below). Water restriction was applied for a maximum of 5 days per week, and on the remaining two (or more) days the rats received free access to water. It was ensured that each rat would receive at least 25 ml of water per 1000g of body weight in a 24-hour cycle (Schellinck et al., 1993; Histed et al., 2012). The rats could continue to perform the experiment for as long as they were motivated to do so, beyond the minimum water intake. At the end of each experimental session water intake was measured, and each rat was supplemented individually, well beyond their daily minimum. The rats were weighed twice a week to ensure stable body weight (at the time of testing their average weight was 527g), and their health was continuously monitored. One rat developed a growth partway through the experiment and was therefore removed from the rest of the experiment (leaving 7, who completed all the conditions).

### Experimental setup

For this experiment we built a custom multisensory motion platform for rats using two individually controllable robotic arms, each with 6 degrees-of-freedom (Fig. 1A; see Suppl. Fig. 1 for photographs of the system). A flat platform was mounted on Robot #1 (Yaskawa Motoman MH5LF) and a chamber, in which the rat was placed, was mounted on the platform (Suppl. Fig. 1B). A second robot (Robot #2, Yasakawa Motoman MH12) held a 90 cm diameter black sphere custom made from fiberglass and lined inside with black acoustic (sound-absorbing) foam. The two robots’ motions could be coupled (i.e. moved together) or decoupled (i.e., each robot moved independently). They were controlled by a master-slave configuration (Yaskawa FS100) to ensure synchronous motions.

The rat was placed in the chamber while the system was in the “disengaged” position – namely, the platform was located outside the sphere (Suppl. Fig. 1A). Then, the system was “engaged” – this moved the platform (via Robot #1) inside the sphere (Suppl. Fig. 1C). In the engaged position, the rat’s head (when protruded to initiate a trial, further details below) was located at the center of the sphere. This was the starting point from which all trials began and to which the system returned at the end of each trial. A flap made from black acoustic foam was used to flexibly close the opening at the back of the sphere, around Robot #1’s arm. This was then covered on the outside with a light-blocking curtain. In this state, the rat was totally encompassed by the sphere environment. A strip of individually programmable LEDs (density 144/m) lined the internal circumference of the sphere (Suppl. Fig. 1D). The LED strip covered 320° of the horizontal circumference around the rat (the remaining 40° was the opening at the back of the sphere through which Robot #1 entered). The LEDs were used to control visual (optic flow) cues for relative motion stimuli (described further in subsection **Experimental conditions**, below).

The rat chamber (made from transparent Perspex) had an opening in the front, which enabled the rats to extend their heads out, to reach three water ports – one located in the center, one to the right and one to the left (Suppl. Fig. 1B). A video camera monitored the real-time position of the rat’s head using video-tracking software (Ethovision XT 12, Noldus). Infrared (940 nm) LEDs were placed beneath the platform. This created a silhouette of the rat’s head for the camera (Fig. 1D) so that video tracking would work in the dark. Water flow in the reward ports was controlled by three separate solenoids (Parker, series 3). Two small speakers were located in the right and the left reward ports (Suppl. Fig. 1B) for training and timing signals.

### Self-motion stimuli

All self-motion stimuli were in the horizontal plane. The heading straight-ahead was defined as 0°. Headings to the right and left were defined by positive and negative angles (relative to straight ahead) respectively (Fig. 1C). Each self-motion stimulus followed a straight (linear) path in a specific heading direction. Each motion stimulus covered 8 cm distance and lasted 1 s. The motion stimulus velocity followed a Gaussian profile, with peak velocity = 0.19 m/s (reached halfway through the stimulus) and peak acceleration = 0.70 m/s^2^ (Fig. 1B).

Inertial self-motion stimuli were generated by moving the motion platform (via Robot #1) in darkness (LEDs off). To avoid any other possible relative motion cues that could arise between the rat and the environment, the sphere was kept at a fixed distance from the rat throughout the motion by coupling the motion of Robot #2 together with Robot #1. Hence, for inertial stimuli the platform and the sphere moved together (Fig. 1A, top panel). This turned out to be an important aspect of the experiment design (because indeed non-visual cues were used for perception of relative self-motions).

Relative self-motion stimuli were generated by displacing the sphere environment (via Robot #2) while the platform remained in place (Robot #1 did not move). Here, Robot #2 moved the sphere to elicit the same relative motion cues that would have arisen from inertial self-motion through a static environment. Accordingly, the sphere was moved in the opposite direction (180°) compared to the direction in which the platform would have been moved, had it been an inertial self-motion stimulus (Fig. 1A, middle panel).

Combined (inertial-relative) self-motion stimuli were generated by displacing the platform (via Robot #1) while keeping the sphere static in world coordinates (no motion of Robot #2; Fig. 1A, bottom panel). In this way, relative motion cues were elicited naturally from the environment, which remained static, as the rat was displaced. This reflects combined cues with no cue discrepancy (Δ = 0°). Combined cue trials were also run with a discrepancy between the inertial and relative motion headings. This is needed in order to measure cue integration weights (Fetsch et al., 2009; Yakubovich et al., 2020). A heading discrepancy of Δ was generated by offsetting both the inertial and relative headings (each by half of Δ) to either side of the trial’s defined heading. Two counterbalanced conditions were run: Δ^+^ = +20° and Δ^−^ = –20°. By convention, positive Δ reflects inertial and relative headings offset to the left and right, respectively, and vice versa for negative Δ. Accordingly, on a trial with heading *h* and discrepancy Δ, Robot #1 generated an inertial motion stimulus with heading *h* – Δ/2, while Robot #2 made a small motion perpendicular to *h* to yield a relative motion stimulus with heading *h* + Δ/2 (according to the net vector of the two Robot’s motions). The “correct” heading for these conditions (for reward purposes etc.) was defined by *h*.

### Heading discrimination task

The rat initiated a trial by poking its head out the chamber, into the zone behind the center reward port (Figure 1D, center). If the rat did not keep its head in this center position throughout the stimulus, the trial was aborted (with no reward). After experiencing a self-motion stimulus (inertial, relative or combined) the rat received a small drop of water (∼5 ul) from the center port. It was then required to report whether the stimulus was to the left or right by turning its head to the left or right water port (detected by video tracking) where it received a bigger reward (∼20 ul) for making a correct choice. The rats were tested one at a time, and monitored by the experimenter. The session was terminated when the rat stopped initiating trials, usually after ∼400 trials (which took ∼40 minutes).

### Training

Rat training took ∼3 months, and comprised 5 main stages: (1) initial handling and exposure to the experimental setup. (2) Gradual exposure to self-motion stimuli. The rats were trained using combined cue stimuli with no cue discrepancy (Δ = 0°). They were trained to maintain their heads in the center detection zone during the stimulus – beginning with weak and short duration stimuli, gradually increasing to 8 cm in 1 s. (3) Heading discrimination training using large headings (±60°). Initially, the reward was given automatically from the left or right reward port, according to the stimulus heading. Reward was then made contingent on the rat making the correct choice. (4) Inertial and relative trials were introduced. The rats immediately performed relative trials with high proficiency, and required extra training for inertial trials. (5) Intermediate (i.e. more difficult) headings were introduced. The rats quickly generalized the task across headings (on the first session these were applied).

### Experimental conditions

#### Baseline

baseline testing began right after training. Four stimulus types were tested: inertial, relative, and two combined (Δ^+^ and Δ^−^) cue stimuli. For each stimulus type, the following headings were tested: ±60°, ±30°, ±15°, and ±7.5°. Stimuli and headings were interleaved pseudo-randomly, according to the method of constant stimuli. Inertial and combined cues were run with LEDs on – namely, each individual LED was turned on with *p* = 0.1, such that a random combination of ∼10% of the LEDs were activated on each trial. This (different combinations of LEDs for each trial) was to mitigate the possibility of the rats using visual cues other than optic flow to solve the task. For the *Baseline* condition 49,365 trials (total) were obtained from the eight rats (> 765 trials per stimulus type, per rat).

#### Baseline (dark)

a few weeks into *Baseline* testing, another stimulus type was added (interleaved with the other four from the *Baseline* condition) – relative self-motion cue in the dark (7,272 trials from 8 rats). Darkness inside the sphere (LEDs off) was verified by measuring luminance (0.002 CD/m^2^) using a photometer (Konica Minolta LS-110). This condition was to compare performance for relative cues without vs. with visual optic flow. Because we found that performance for relative cues was only marginally worse in the dark vs. light (Fig. 3C) a subsequent set of control conditions (described below) were run, to identify what sensory cues the rats primarily used for perceiving relative stimuli.

#### No Sphere

to verify that the rats indeed used relative motions of the sphere to perceive relative stimuli (vs. other cues, such as noises from the robots) we dismounted the sphere, and reran the experiment with the four standard stimulus types (like *Baseline*): inertial, relative, and two combined (Δ^+^ and Δ^−^). The room had ambient light, hence there was no ‘dark’ stimulus type. In this condition the robots performed the identical motions as before (like *Baseline*) however Robot #2 had nothing attached, and therefore just made motions above the head of the rat. Thus, the rat was exposed to the environment of the room during testing. This condition comprised 9,615 trials from 7 rats.

#### Fan

to create turbulence that would interfere with the airflow during self-motion cues, we mounted a 8×8 cm computer fan (Delta electronics) above the chamber, facing forward. The fan was turned on for the whole duration of the block. Five stimulus types were tested – the four standard stimulus types (like *Baseline*) as well as relative self-motion in the dark. This condition comprised 43,267 trials from 8 rats.

#### Fan Noise

to confirm that fan noise did not damage performance, we ran a control condition in which we covered the fan with a Perspex box to block airflow but not fan noise. The fan was turned on for the whole duration of the block. Five stimulus types were tested – the four standard stimulus types (like *Baseline*) as well as relative self-motion in the dark (27,763 trials from 7 rats).

#### Windshield

to block airflow passively (without a fan) we positioned a half-sphere windshield custom-made from clear Perspex (by cutting a 23cm diameter lamp shade in half) in front of the chamber, on the far side of the reward ports. This blocked airflow mainly from the front (but could allow some airflow from the sides). Five stimulus types were tested – the four standard stimulus types (like *Baseline*) as well as relative self-motion in the dark (27,659 trials from 7 rats).

#### Whiskers Cut

for this condition we trimmed all the large whiskers (macro vibrissae) of the rats. This procedure was performed under isoflurane anesthesia. All facial whiskers, on both sides of the snout (rows A-E, arcs 0-6), the 3 trident whiskers located on the chin (The et al., 2013; Chorev et al., 2016), as well as long whiskers located above the eyes and other visible whiskers were trimmed, to <1mm length. This was repeated, if needed, after 2-3 days. Five stimulus types were tested – the four standard stimulus types (like *Baseline*) as well as relative self-motion in the dark (29,389 trials from 7 rats).

### Airflow measurement

An airspeed meter (ATAL, AT-HD29371TC2-5M) was positioned near the opening of the behavioral chamber (without the rat) to measure airflow during the self-motion stimuli (Suppl. Fig. 4). Airflow measurements were similar for relative and combined cues (red and green, respectively) and close to zero for the inertial cue (blue). This indicates that similar airflow was elicited for relative self-motion stimuli – both when elicited from inertial motion of the rat (in combined conditions) and when elicited by motion of the sphere (in relative conditions). Furthermore, airflow was canceled out in inertial stimuli. The airflow measures were not intended to quantify exact airflow timing (due to technical limitations of the apparatus, e.g. a slow return to pre-stimulus baseline). Rather they were just intended to show that relative airflow resulting from relative or combined stimulus types were comparable (and that this was absent for inertial cues).

### Data acquisition and analysis

The experiment and robots were controlled by custom software written in C#. Data analyses were performed using Matlab 2013b (MathWorks). Several measures were used to quantify performance. First, percent correct was used, as a model-free measure of performance, for each stimulus type, condition and rat. To be comparable across stimulus types, conditions and rats (even if the number of trials could slightly differ per heading) percent correct was first calculated per heading, and then averaged over headings (all stimulus types, conditions and rats had the same headings). Although the cue discrepancy could reduce percent correct for combined cues (the Δ could lead to a PSE bias), performance was still shown to be significantly better in combined cue conditions, and therefore, this was not an issue.

Psychometric curves (cumulative Gaussian) were fit to the proportion of rightward choices as a function of heading (per stimulus type, condition and rat) using the Psignifit-4 toolbox for Matlab (Schütt et al., 2016). We extracted three psychometric parameters: the point of subjective equality (PSE), defined by the mean (μ) of the Gaussian fit, the behavioral threshold, defined by the standard deviation (SD) of the Gaussian fit, and the lapse rate, defined by the asymptotes of the fit (assumed symmetrical for right and left; Shalom-Sperber et al., 2022). Priors for the psychometric fit parameters were set by defining the stimulus range as [-90°, 90°] in Psignifit. This resulted in: i) a flat PSE prior within this range (with decreasing probability outside this range until ±180°), and ii) a flat threshold prior up to 55° (with decreasing probability outside this range until 164°). iii) For lapse rates, we set a flat prior within the range 0 to 0.15, with decreasing probability outside this range until 0.3 (lapse rates ≥ 0.3 were not allowed, to prevent overfitting/ underestimating thresholds). The goodness-of-fit of the psychometric functions was evaluated using pseudo-*R²* (Hosmer & Lemeshow, 1989).

The empirical (observed) combined cue thresholds (𝜎_𝑐𝑜𝑚𝑏𝑖𝑛𝑒𝑑_𝑜𝑏𝑠_) were calculated (per rat, and condition) by the geometric mean of the psychometric fit thresholds from the two combined cue conditions (Δ^+^ and Δ^−^). These (observed) thresholds were compared to the Bayesian-predicted combined cue thresholds, calculated from the inertial and relative psychometric fit thresholds (per rat and condition), as follows (Angelaki et al., 2009):

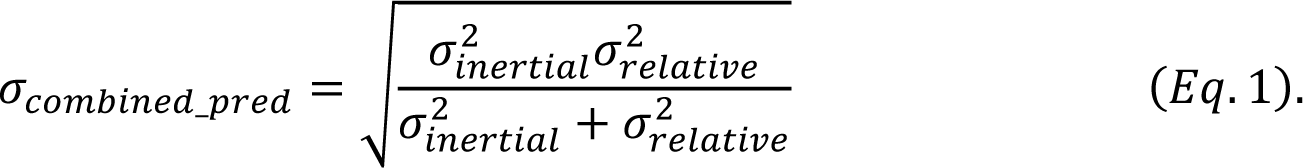

The empirical (observed) weights for inertial and relative motion cues were estimated from the PSE fits of the two combined cue conditions as follows (Fetsch et al., 2012; Yakubovich et al., 2020):

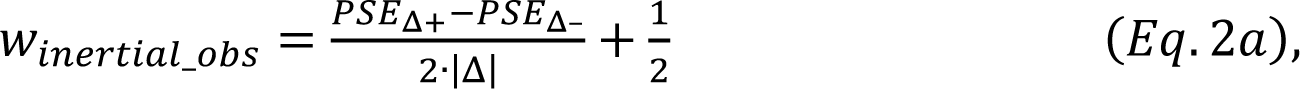

and

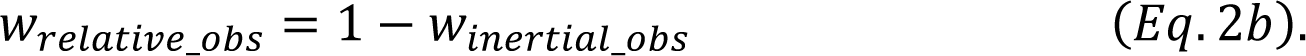

These (observed) weights were compared to the Bayesian-predicted weights, which were calculated from the inertial and relative psychometric fit thresholds (per rat and condition), as follows:

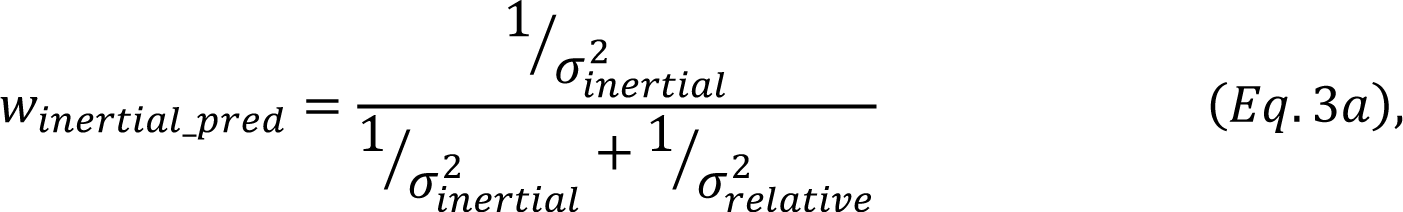

and

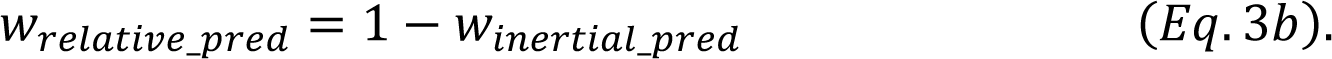

### Statistical analysis

Statistical analyses were performed using JASP (JASP Team, 2018). To compare between stimulus types (inertial, relative and combined) in the *Baseline* condition we used repeated measures ANOVA and accordingly, Holm post-hoc comparisons. To test the effects of the control conditions, performance was compared vs. *Baseline* (for inertial and relative cues). Similarly, performance in the control (dark) conditions was compared vs. *Baseline (dark)*. These comparisons were generally done using one-tailed paired *t*-tests and Bayes Factors, with the alternative hypothesis (H_1_) that performance in the control condition was worse. This was based on the rationale that the control condition aimed to impair performance. An exception to this was the *No-Sphere* condition. Here, we expected performance for the ‘inertial’ stimulus to possibly be better than *Baseline* because, without the sphere, the external room naturally provided relative motion cues (it thus essentially became like a combined stimulus). Hence, comparisons using ‘inertial’ and combined stimulus types in the *No-Sphere* condition were two-tailed.

Bonferrroni correction was used to correct for multiple comparisons, as follows. For the inertial cue (percent correct comparisons, Tables 1 and 2B) five control conditions (*No Sphere*, *Fan*, *Fan noise*, *Windshield*, *Whiskers cut*) were compared to *Baseline*. Thus, the resulting *p*-values were multiplied by five. For the relative cue (percent correct comparisons, Tables 1 and 2A) six control conditions with LEDs on (*No Sphere*, *Fan*, *Fan noise*, *Windshield*, *Whiskers cut*, *Baseline (dark)*) were compared to *Baseline*, and five control conditions with LEDs off (*Fan (dark)*, *Fan noise (dark)*, *Windshield (dark)*, *Whiskers cut (dark)*) were compared to *Baseline (dark)*. Thus, the resulting *p*-values were multiplied by six and five, respectively. For the combined cue (percent correct comparisons, Table 1) two comparisons were made, thus the resulting *p*-values were multiplied by two. Threshold comparisons (Table 3) were confirmatory of the percent correct comparisons (not independent hypotheses) and thus followed the same rationale of corrections presented above, but with one less analysis (thresholds were not calculated for the *No Sphere* condition because the drop in relative cue performance was too extreme to fit a psychometric curve).

Correlations between the observed and Bayesian-predicted combined cue thresholds, and between the observed and Bayesian-predicted integration weights were used to test for cue reweighting (pooling across rats and conditions). Bayes Factors (for these correlations and the comparisons in Tables 1-3) with values *BF_10_* > 3 or *BF_10_* < ⅓ were considered substantial evidence for H_1_ or H_0_, respectively (Jarosz & Wiley, 2014). Otherwise: 1 > *BF_10_* > 3 or ⅓ < *BF_10_* < 1 were considered anecdotal evidence (for H_1_ or H_0_, respectively) or inconclusive (*BF_10_* ∼1).

### Data and code availability

The data and analysis scripts (Matlab) for this study can be found at: https://osf.io/n8uds/?view_only=68b3ce1325274f58bac36582fec68105.

## Acknowledgements

We would like to thank Avraham Elkaras for programming, Judith Sonn for management and David Swissa for mechanical assistance. We thank Orly Halperin, Avigail Spolter and Elad Goldberg for help with animal training and testing. This research was supported by grants from the Israeli Centers of Research Excellence (I-CORE, Center No. 51/11) and the Israel Science Foundation (ISF, grant No. 1291/20) to AZ.

## Authors’ roles

LP performed experiments, analyses, and wrote the paper. TH performed analyses and wrote the paper. AZ designed and supervised the study, and wrote the manuscript.

## Supplementary Material

**Supplementary Figure 1:**
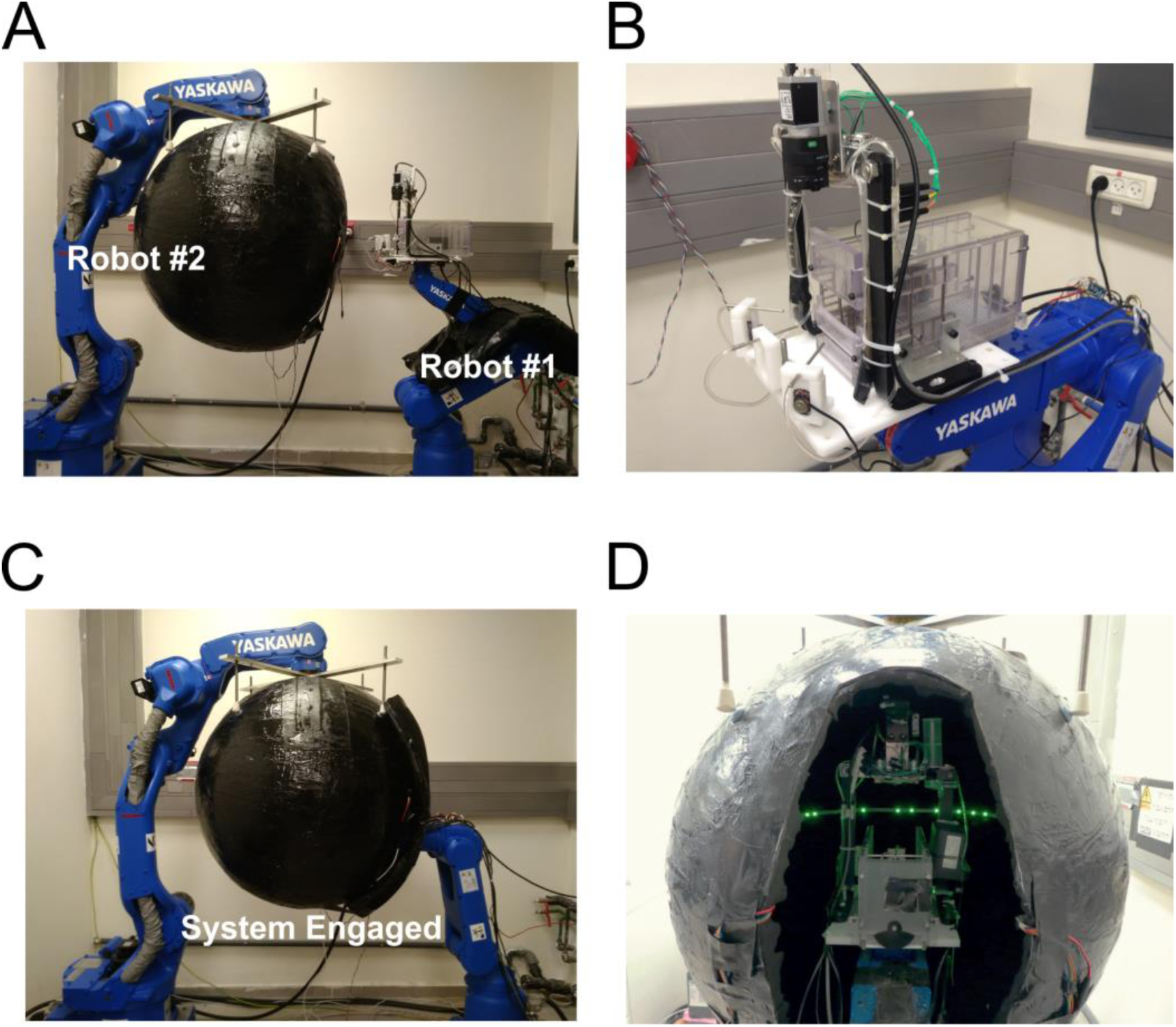
Photographs of the rat motion system. Robot #1 supports and moves the rat platform. Robot #2 supports and moves the black sphere (environment). (A) The motion system ‘disengaged’ (i.e., rat platform not inside the sphere). (B) The rat platform. (C) The motion system ‘engaged’ (i.e., rat platform inside the sphere). (D) View into the sphere from behind Robot #1 (system engaged). Green LEDs line the inside circumference of the sphere. The black flap, which normally closes this opening during experiments (see Panel C), was left open here for visibility into the sphere.

**Supplementary Figure 2:**
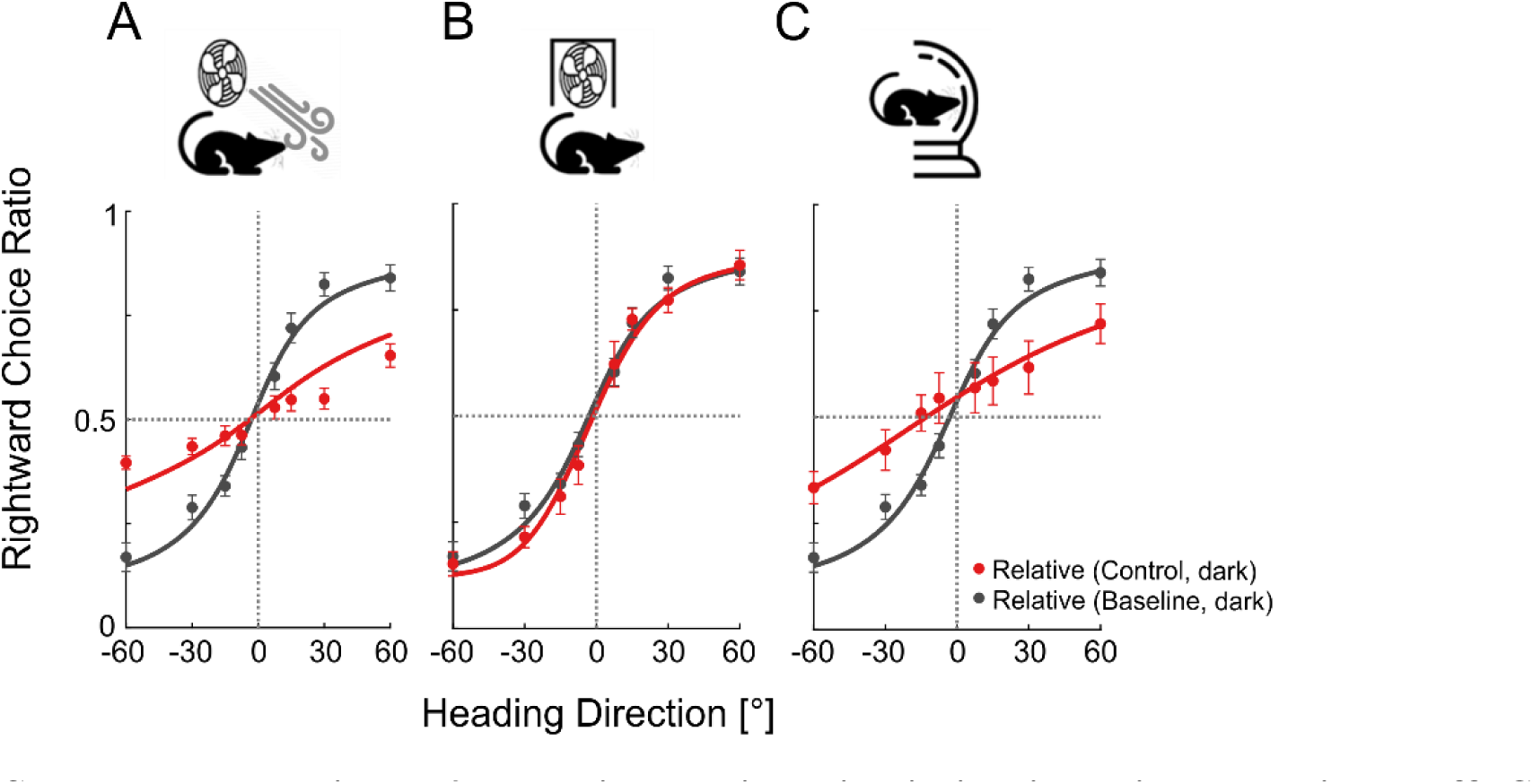
Relative motion discrimination with LED lights off. Conventions are the same as Fig. 4, top row. However, these experiments were run in the dark (the LED lights inside the sphere were off). All pseudo-*R²* values were > 0.87.

**Supplementary Figure 3:**
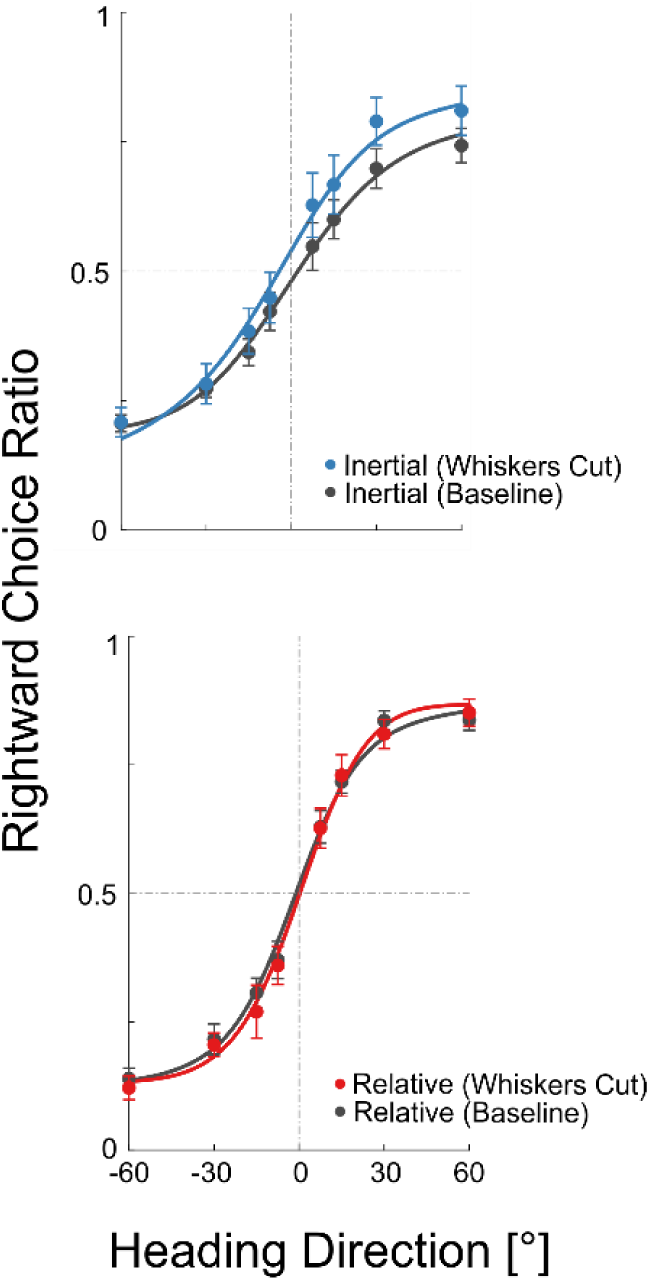
Intact self-motion perception with whiskers cut. Conventions are the same as Fig. 4, but here with data from the *Whiskers Cut* (control) condition. All pseudo-*R²* values were > 0.97.

**Supplementary Figure 4:**
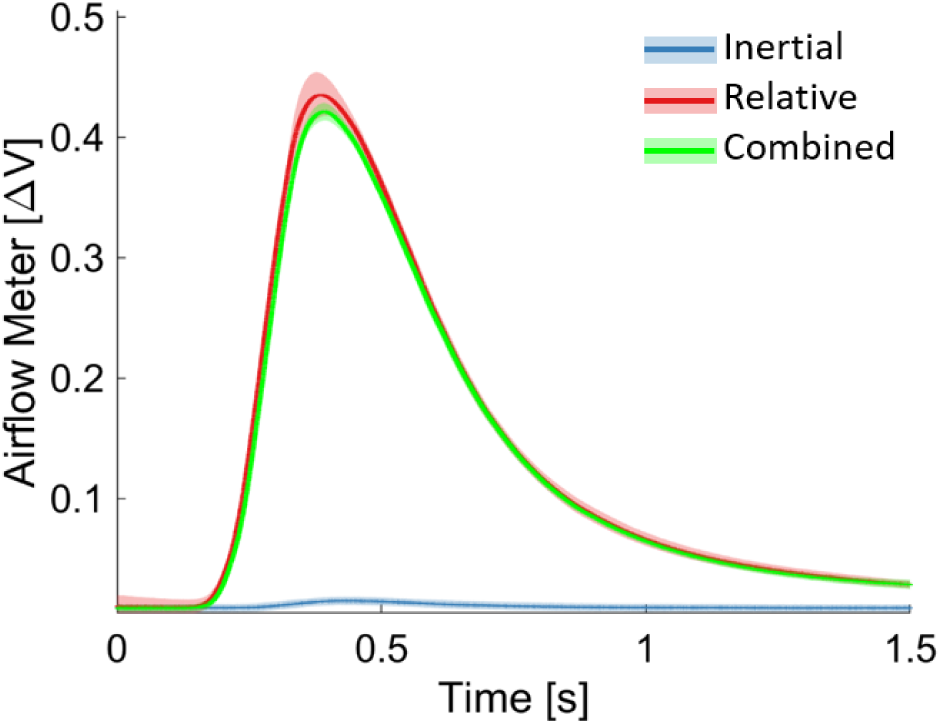
Airflow measurement. An airflow meter, positioned near the opening of the behavioral chamber, measured the airflow generated by the stimuli (20 repetitions each, at 0° heading). Solid lines and shaded regions mark the mean ± SD, per stimulus. Measurements (change in airflow meter voltage from baseline; ΔV) were similar for relative and combined motion cues (red and green, respectively) and close to zero for inertial motion (blue).

